# Multiheme selenoenzyme essential for elemental sulfur respiration

**DOI:** 10.1101/2025.11.18.688581

**Authors:** Hisaaki Mihara, Takuya Yoshizawa, Yukiko Izu, Wanjiao Zhang, Masao Inoue, Nana Shimamoto, Ryuta Tobe, Riku Aono, Tatsuo Kurihara, Hiroyoshi Matsumura

## Abstract

Microbial reduction of elemental sulfur is a pivotal process in the global sulfur cycle^1^. Insoluble elemental sulfur can serve as a terminal electron acceptor during anaerobic respiration, producing sulfide^2,3^. However, the molecular mechanisms underlying sulfur reduction remain poorly understood. Here we show that an evolutionarily conserved multiheme cytochrome *c* selenoprotein (MccSep) is essential for elemental sulfur reduction in a sulfur-respiring bacterium. Our biochemical and structural analyses revealed that MccSep comprises four identical subunits, each containing five *c*-type hemes and one selenocysteine residue (U325). The enzyme catalyzes polysulfide reduction at a unique active site, where C239 coordinates with the heme iron center, while U325 plays a crucial role in the catalytic process. Genetic analysis further revealed the critical roles of U325 and C239 in driving *in vivo* elemental sulfur respiration. These findings unveiled a selenium-sulfur-dependent catalytic chemistry on a heme center during polysulfide reduction, expanding the scope of biological catalysis and deepening our molecular understanding of bacterial sulfur respiration.

## Main

Elemental sulfur (S^0^) is widespread in the environment and originates from volcanoes, hot springs, hydrothermal vents, and microbial sulfur metabolisms^4^. Despite its low solubility in water^5^, S^0^ serves as a terminal electron acceptor in microbial anaerobic respiration, producing sulfide (H_2_S/HS^-^) and playing a pivotal role in the biogeochemical sulfur cycle^1,2^. The dissimilative S^0^-reducing bacterium—*Geobacter sulfurreducens* PCA—belongs to the phylum Desulfobacterota^6^, which also includes *Desulfuromonas acetoxidans*—the first S^0^-reducing bacterium discovered in 1976^7^. Although these bacteria reduce S^0^, they cannot use oxidized sulfur compounds, such as sulfate, thiosulfate, and sulfite, for anaerobic respiration^7,8^. Despite decades of research^9,10^, the molecular basis of sulfur reduction in these bacteria has remained elusive for nearly half a century.

*G. sulfurreducens* and *D. acetoxidans* possess numerous multiheme cytochrome *c* (MCC) proteins (120 and 50, respectively), which have been postulated to function as extracellular electron transfer conduits, electron carriers in the periplasm, and terminal oxidases^11,12^. The structures of MCC enzymes from different bacteria revealed that catalysis occurs either via direct axial coordination of a substrate to the active-site heme iron, as observed in MCC nitrite reductase (NiR)^13^ and sulfite reductase (MccA)^14^, or via a catalytic Cys residue involved in His/Cys-ligation to heme *c*, as observed in thiosulfate oxidase (TsdA)^15,16^.

We previously found an uncharacterized periplasmic MCC protein^17^, encoded by *gsu2937*–*2936* (also termed *extKL*^18^) in the *G. sulfurreducens* PCA genome that may contain a selenocysteine (Sec) residue, the 21^st^ amino acid encoded by the UGA codon, which typically serves as a stop codon^19^. This putative MCC selenoprotein, designated MccSep, was predicted to possess five *c*-type heme-binding motifs (CxxCH) and a Sec residue^17^ (Fig. 1a). Sec-containing enzymes catalyze crucial redox reactions via the high reactivity of Sec, which plays a role in bacterial anaerobic energy metabolisms^20^. Furthermore, our recent transcriptome analyses had revealed upregulation of *gsu2937*–*2936* (*mccsep*) expression in response to S^0^ supplementation^21^, suggesting its possible role in sulfur metabolism. The electron transfer capabilities of MCCs, the catalytic properties of Sec, and sulfur-responsive expression patterns prompted us to hypothesize that MccSep functions as an oxidoreductase in sulfur reduction. Here, we demonstrate that MccSep is a key polysulfide reductase for S^0^ respiration in *G. sulfurreducens* PCA. Genes encoding MccSep homologs are widespread across bacterial genomes of potential S^0^ reducers, including numerous metagenomes. Using a combination of biochemical, crystallographic, and genetic analyses, we demonstrate that MccSep catalyzes polysulfide reduction via a Sec-, Cys-, and heme-dependent mechanism, thereby providing a molecular framework for understanding bacterial S^0^ respiration.

**Fig. 1.**
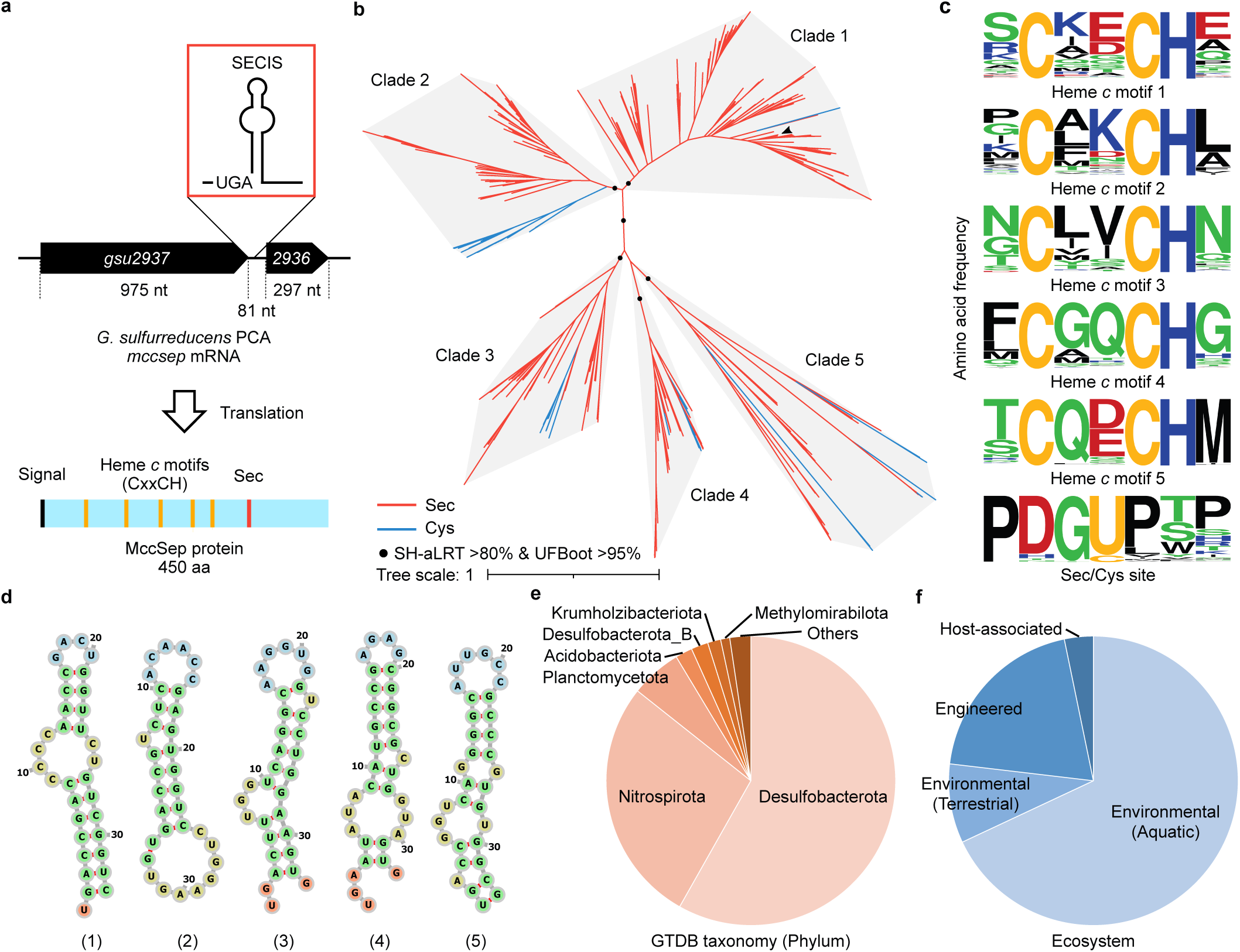
Bioinformatics analysis of MccSep homologs. **a**, Schematic representation of the *gsu2937-2936* mRNA and its gene product, MccSep, from *G. sulfurreducens* PCA. The diagram shows the UGA codon corresponding to Sec and the downstream SECIS-like element, which is a predicted mRNA stem–loop structure required for Sec insertion during translation. MccSep schematic highlights the signal peptide, heme-binding sites, and the Sec residue. **b**, Unrooted maximum-likelihood phylogenetic tree of MccSep homologs harboring a conserved Sec/Cys residue. The tree was constructed from a multiple sequence alignment of the centroid sequences representing 455 clusters at 90% amino acid sequence identity. Branches containing Sec-and Cys-type MccSep homologs are colored red and blue, respectively. Major clades (1–5) are shaded gray. Black circles indicate branches separating major clades with bootstrap values of SH-aLRT >0.80 and UFBoot >0.95. **c**, Amino acid frequency profiles of five conserved heme *c*-binding motifs and the Sec/Cys site, visualized using WebLogo. **d**, Predicted RNA secondary structures of SECIS-like elements from Sec-containing representatives of each major clade, generated using MXFold2. The analyzed sequences include the UGA codon and 30 nt downstream from the following representative locus tags: 1, *Geobacter sulfurreducens* (GS_RS14775); 2, *Desulfurivibrio alkaliphilus* (DAAHT2_RS12195); 3, *Thermosulfurimonas dismutans* (TDIS_RS01570); 4, Nitrospirota bacterium (ENH01_02100); 5, *Candidatus* Brocadia sinica (BROSI_RS02360). **e**, Taxonomic distribution of 684 genomes encoding MccSep homologs, classified at the phylum level using GTDB-Tk. **f**, Ecosystem distribution of 588 metagenome-assembled/single-amplified genomes harboring MccSep homologs.

## Distribution of MccSep homologs

To demonstrate that MccSep represents a new bacterial selenoprotein family, we first performed bioinformatics analysis of its sequence characteristics and genomic distribution. A total of 637 non-redundant MccSep homologous sequences containing the conserved Sec/Cys residue were retrieved from the NCBI genome database, a subset of which (394 sequences) was recovered using UGA stop codon read-through analysis (Supplementary Datasets 1,2). Phylogenetic analysis revealed that the MccSep homologs formed a protein family consisting of five major clades (Fig. 1b). The Cys-type counterparts emerged from the clade of Sec-type sequences, similar to those previously described for other selenoproteins^22^. All protein sequences for phylogenetic analysis showed five conserved heme *c*-binding motifs and Sec/Cys sites as unique characteristics of the MccSep family (Fig. 1c). In Sec-type representatives from each clade, mRNA segments containing UGA and its 30 nt downstream sequence were predicted to form Sec insertion sequence (SECIS) element-like stem–loop structures^19^ (Fig. 1d). Genome taxonomy annotation revealed that nearly all MccSep family sequences originated from bacteria, primarily from the metagenome-assembled genomes of uncultured taxa (Fig. 1e and Supplementary Datasets 1,2). MccSep family sequences were mainly distributed in the phyla Desulfobacterota (including the genus *Geobacter*) and Nitrospirota (mainly in the class Thermodesulfovibrionia). These taxa are known for their diverse capabilities in sulfur reduction or disproportionation^8,23,24^. Most MccSep-encoding metagenome-assembled genomes are derived from aquatic environments, such as marine, freshwater, groundwater, and hydrothermal vents, suggesting their potential role in these habitats (Fig. 1f and Supplementary Datasets 1,2).

## Identification of MccSep

To confirm the production of Sec-type MccSep by *G. sulfurreducens* PCA, we used an anti-GSU2936 antibody generated against the 11-kDa GSU2936 protein, which was recombinantly expressed in *Escherichia coli* BL21(DE3). Western blot analysis of the crude extracts of *G. sulfurreducens* PCA revealed a single band at approximately 49 kDa, close to the molecular mass of the predicted *mccsep* product (Fig. 2a), suggesting that the UGA codon located in-frame between *gsu2937* and *gsu2936* underwent read-through, leading to the production of a single polypeptide, MccSep (Fig. 1a). Western blot analysis of the bacterial lysate fractions indicated the localization of MccSep in the periplasm (Fig. 2b). Consistently, 97% of MccSep homologs are predicted by DeepLocPro^25^ to possess periplasmic localization signals (Supplementary Datasets 1,2).

**Fig. 2.**
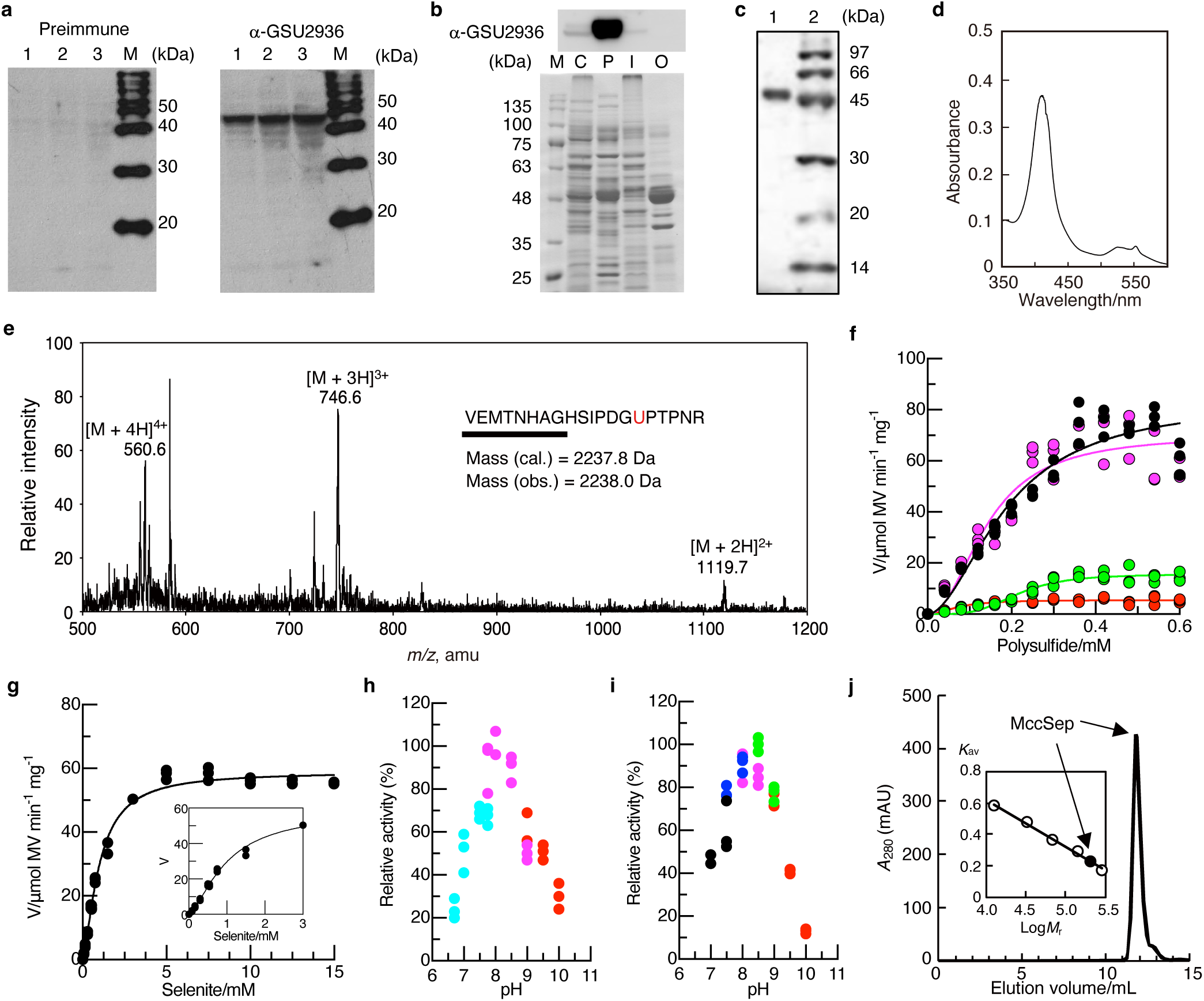
Characterization of MccSep. **a**, Western blotting of MccSep in the crude extract of *G. sulfurreducens* PCA using a preimmune antisera (left) or anti-GSU2936 antibody (right). Protein loaded were 9.1 µg (lanes 1), 14 µg (lanes 2), and 18 µg (lanes 3). MagicMark XP Western Protein Standard (Thermo Fisher Scientific) was loaded onto lanes M. **b**, Subcellular fractions (cytoplasm, C; periplasm, P; inner membrane, I; outer membrane, O) of *G. sulfurreducens* were analyzed by western blotting using anti-GSU2936 (upper panel). The lower panel shows replicate gels visualized by CBB staining, indicating the separation of the subcellular fractions and the amounts of proteins loaded. Equal amounts (12 µg) of total protein were applied to each fraction. **c**, SDS-PAGE of MccSep (0.5 µg) purified from *G. sulfurreducens* PCA. The gel was stained with CBB. **d**, UV–vis absorption spectrum of dithionite-reduced MccSep. MccSep (0.34 mg/mL) was reduced with 19 mM dithionite in 20 mM Tris-HCl (pH 7.8). **e**, ESI–MS analysis of the Sec-containing tryptic peptide of iodoacetamide-alkylated MccSep. The positive ion mass spectrum of the purified peptide is shown. The underlined sequence agrees with that determined by Edman sequence analysis. **f**, **g**, Steady-state kinetic analysis of MccSep WT (black), R83A (green), C240A (magenta), and K372A (red) with polysulfide (**f**) or selenite (**g**) as a substrate. The lines are the best fit to an allosteric sigmoidal equation. The data shown are from experiments performed in triplicate. Polysulfides were a mixture of different chain lengths, and their concentration is expressed as the total sulfur equivalents. **h**, **i**, Effect of pH on MccSep activity with polysulfide (**h**) was determined at varying pH using 50 mM each of phosphate (cyan), Bicine-NaOH (magenta), and glycine-NaOH (red) buffers. Effect of pH on MccSep activity with selenite (**i**) was determined using 50 mM each of MOPS-NaOH (black), HEPES-NaOH (blue), Bicine-NaOH (magenta), glycylglycine-NaOH (green), and glycine-NaOH (red) buffers. The data shown are from experiments performed in triplicate. **j**, Size exclusion chromatography analysis of MccSep using a Superdex 200 Increase 10/300 GL column. The inset shows the calibration curve of the column using a mixture of standard marker proteins indicated by white circles.

Native MccSep was purified from the crude extract of *G. sulfurreducens* PCA using sequential column chromatography (Fig. 2c). Edman sequencing of purified MccSep identified an N-terminal peptide (EKAGIGWQETIVAKS) corresponding to residues 26–40 of the full-length protein (Extended Data Fig. 1), consistent with the predicted cleavage of its signal peptide (residues 1–25) for periplasmic localization (Fig. 2b). Purified MccSep was red, consistent with the predicted heme cofactors. UV–vis absorption spectrum of dithionite-reduced MccSep exhibited absorption maxima at 420 nm (γ), 523 nm (β), and 552 nm (α), which are characteristic of cytochrome *c* proteins^10^ (Fig. 2d). Purified MccSep contained 4.9 ± 0.1 iron atoms per subunit, determined colorimetrically using the iron-bathophenanthroline disulfonate complex formation^26^, consistent with the prediction that MccSep has five heme *c* cofactors per subunit. The presence of Sec in MccSep was confirmed by fluorometric determination of selenium^27^ and by ESI–MS analysis of tryptic fragments derived from iodoacetamide-alkylated MccSep. Fluorometric analysis revealed 0.96 ± 0.13 selenium atoms per subunit. In ESI–MS analysis, the observed molecular mass of the purified Sec-containing peptide (VEMTNHAGHSIPDG<u>U</u>PTPNR) with *Se*-carbamidomethylated Sec (2238.0 Da) closely matched the calculated value (2237.8 Da) (Fig. 2e). These results demonstrate the in-frame incorporation of Sec at the UGA codon in the *mccsep* gene, yielding the MccSep protein.

## MccSep is a polysulfide reductase

To further characterize MccSep, we purified recombinant MccSep carrying a C-terminal Strep-tag, expressed in an *mccsep*-deficient (Δ*mccsep*) strain of *G. sulfurreducens* PCA harboring the expression vector pMcasMccSepStag (Extended Data Figs. 2 and 3a,b). The UV–vis absorption spectrum of dithionite-reduced recombinant MccSep closely resembled that of the native MccSep (Extended Data Fig. 3c and Fig. 2d). We next examined MccSep activity toward polysulfide, based on the hypothesis that it functions as a periplasmic enzyme involved in sulfur reduction. In addition, we assayed other potential substrates, including those previously reported for MCC enzymes, using dithionite-reduced methyl viologen (MV) as the electron donor. Consistent with this hypothesis, MccSep efficiently catalyzed the reduction of polysulfide and, to a lesser extent, selenite (Figs. 2f,g). MccSep-catalyzed polysulfide reduction followed allosteric sigmoidal kinetics, with a *V*_max_ of 83 ± 7 μmol MV oxidized min^-1^ mg^-1^, *K*_0.5_ of 0.19 ± 0.02 mM, and a Hill coefficient of 1.9 ± 0.3 (Fig. 2f and Extended Table 1). Selenite reduction showed similar kinetics but with lower efficiency (*V*_max_ of 59 ± 1 μmol min^-1^ mg^-1^, *K*_0.5_ of 0.99 ± 0.04 mM, and a Hill coefficient of 1.5 ± 0.1) (Fig. 2g). The lower *K*_0.5_ (fivefold) and higher *V*_max_/*K*_0.5_ (sevenfold) for polysulfide clearly indicate that it is the preferred substrate of MccSep. The enzyme activity for polysulfide and selenite was optimum at pH 8.0–8.5 (Figs. 2h,i). In contrast, no detectable activity was observed with thiosulfate, tetrathionate, sulfite, sulfate, nitrite, nitrate, selenate, tellurate, or arsenite (Supplementary Data 1), none of which are known terminal electron acceptors supporting the growth of *G. sulfurreducens* PCA^8^.

## Crystal structures of MccSep

To investigate the structural basis of the function of MccSep, we determined its crystal structure at 1.75 Å resolution (Fig. 3a and Extended Data Table 2). The crystals contained one MccSep tetramer per asymmetric unit, consistent with the homotetrameric structure of 200 kDa in solution, as determined by size-exclusion chromatography analysis (Fig. 2j). Each monomer formed a mushroom-like structure with a cap and stem (Fig. 3b). The N-terminus α-helical cap bound five *c*-type hemes, and the C-terminus folded an immunoglobulin-like (Ig-like) β-sandwich stem^28^ with an extended β-hairpin. Five *c*-type hemes were anchored by CxxCH motifs and numbered 1–5 from the N terminus of the MccSep sequence (Fig. 3b and Extended Data Fig. 1). No remarkable differences in the heme-binding sites were observed among the four monomers. Hemes 2 and 5 branched from heme 4 and formed a Y-shaped arrangement. The distances between neighboring iron atoms of the hemes ranged from 9.5 to 11.8 Å. The Y-shaped arrangement of the five hemes in MccSep was superimposable on that of the core five hemes in other MCC proteins^14,29^, despite their overall protein sequences and structures being unrelated (Extended Data Fig. 4). A similar branched heme configuration in other MCC, such as nitrite reductase, has been proposed to transiently store electrons during turnover^30^. The structure of MccSep further suggests that heme 1, which is exposed to the solvent, likely serves as the initial entry point for electrons from an unidentified electron donor (Fig. 3b). The electrons may then be transferred through hemes 3 and 4, ultimately reaching heme 2 at the active site during catalysis, as observed in other MCC enzymes^13^.

**Fig. 3.**
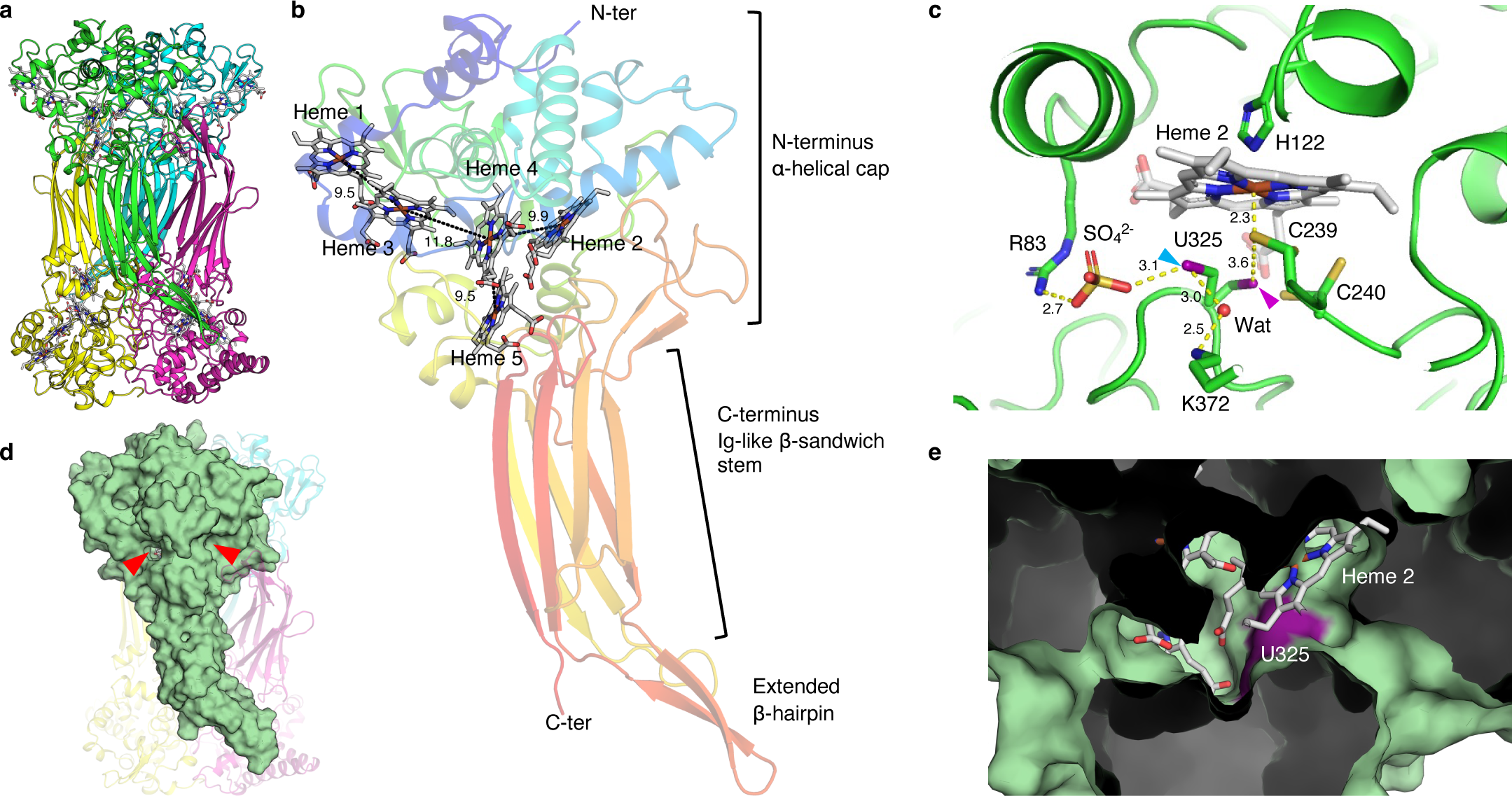
Crystal structure of MccSep. **a**, Side view of tetramer MccSep. Each protomer is represented in a cartoon form with green, magenta, cyan, and yellow. Hemes are shown as grey stick models. **b**, Rainbow-colored cartoons of the protomer structure, from the blue N terminus to the red C terminus. Hemes are shown as stick models. Dashed lines indicate the distances between neighboring iron atoms (Å). **c**, Close-up view of the heme 2 ligand binding site. C239 coordinates the iron atom of heme 2 at a distance of 2.3 Å. U325 in two alternative conformations, conf1 (magenta arrowhead) and conf2 (cyan arrowhead), positioned close to the sulfur atom of C239 at 3.6 Å and the sulfate oxygen atom at 3.1 Å distances, respectively. **d**, Protomer structure of the MccSep tetramer is shown as a green surface model. Red arrowheads indicate edges of the channel. **e**, Cross-section of the channel structure. The position of U325 is colored in purple.

## Active site with Sec and heme-ligating Cys

Among the five heme groups, four (hemes 1, 3, 4, and 5) are coordinated by His residues on both the proximal and distal sides, forming a typical bis-His ligation (Extended Data Fig. 5). In contrast, heme 2 displays an unusual axial ligation by H122 (proximal) and C239 (distal) (Fig. 3c), both of which are strictly conserved among the homologs. This type of His/Cys coordination has previously been identified as a catalytic feature in several sulfur-metabolizing MCC enzymes^15,31^. U325, which extends from the C-terminal Ig-like domain, is positioned near the distal pocket of the His/Cys-ligated heme 2, thereby forming the catalytic center of MccSep (Fig. 3c). The critical roles of U325 and C239 are supported by the severe loss of activity observed in the C239H, U325A, and U325C variants for both polysulfide (Fig. 2f, Extended Data Table 1, and Supplementary Data 2) and selenite reduction (Supplementary Data 3). In contrast, substitution of the adjacent but non-conserved C240 with Ala had little effect on activity (Fig. 2f and Extended Data Table 1). Electron density analysis of the wild-type (WT) crystal structures revealed that the U325 side chain, together with the D323–G324–U325–P326 loop, adopts two alternative conformations, referred to as conformation 1 (conf1; approximately 70% occupancy) and conformation 2 (conf2; approximately 30% occupancy) (Fig. 3c and Extended Data Fig. 6). In comparison, the U325A mutant exhibited a single conformation (Extended Data Table 1 and Extended Data Fig. 6).

## Insights into substrate binding and Sec-mediated catalysis

The MccSep structure features a substrate-accessible channel spanning the N-terminal heme-binding domain and the C-terminal Ig-like domain, extending across the active site (Figs. 3d,e), as observed in other MCC enzymes^32^. At the entrance of this channel, a sulfate ion (an analog of the selenite substrate) is located near U325 in its conf2 conformation, where it interacts with R83 (Fig. 3c). Although a binary complex structure with polysulfide was not obtained, anomalous difference Fourier maps of WT crystals soaked with selenite (SeO_3_^2-^) or selenate (SeO_4_^2-^), as well as of U325A crystals soaked with selenate, show that the sulfate ion bound to R83 is replaced by selenite or selenate (Extended Data Figs. 7a–c). In the WT crystal soaked with selenite, the electron density of the Se_γ_ atom of U325 (conf2) appears continuous with that of the bound selenite (Extended Data Fig. 7a), whereas no such continuity is observed in the structures containing sulfate or selenate, which are non-substrate analogs. These observations suggest that conf2 represents a catalytically relevant state in which U325 nucleophilically attacks the substrate to form a covalent intermediate.

In the WT structure, the Se_γ_ atom of U325 (conf2) forms a water-mediated interaction with the N_ϵ_ atom of K372 (Fig. 3c), although the corresponding electron density for the water molecule is relatively weak, likely due to low occupancy or high mobility. Mutational analyses showed that substitution of K372 with Ala markedly reduced *V*_max_ without significantly affecting *K*_0.5_ (Fig. 2f, Extended Data Table 1, and Supplementary Data 3). A similar decrease in activity was observed for the R83A mutant, consistent with a role of R83 in orienting the substrate and supporting the catalytic function of U325.

## Role of MccSep in S^0^ respiration

We investigated the *in vivo* role of *mccsep* in S^0^ respiration in *G. sulfurreducens* PCA by assessing halo formation on a dispersed S^0^/polysulfide agar medium prepared by adding a polysulfide solution to NBAFYE medium followed by partial air oxidation. Under these conditions, residual polysulfide species remain in equilibrium with finely dispersed S⁰, as previously described for the abiotic conversion of polysulfide to S⁰ under near-neutral, micro-oxic conditions^33,34^. Thus, halo formation reflects the net turnover between sulfur species (S⁰ and Sₙ²⁻) rather than the direct microbial reduction of solid S⁰. Parent *G. sulfurreducens* PCA colonies grown on this medium displayed a distinct clear halo (Fig. 4a), consistent with the conversion of water-insoluble S^0^ into soluble polysulfide via chemical reactions between S^0^ and microbially produced sulfide (Fig. 4b) under anaerobic conditions^35,36^. The Δ*mccsep* strain exhibited a markedly smaller halo than the parent strain (Fig. 4a). However, introducing the *mccsep* expression plasmid (pMcasMccSep) into Δ*mccsep* fully restored the halo to parent-strain levels. The absence of *mccsep* also led to a substantial decrease in sulfide production from S^0^ in liquid culture compared with the parent strain (Fig. 4c). Complementation with variant plasmids expressing U325A, U325C, C239H, R83A, or K372A failed to restore the phenotype (Fig. 4a), confirming that these residues are essential for MccSep function. Furthermore, Δ*mccsep* showed markedly impaired growth and sulfide production in a minimum medium supplemented with S^0^ as the sole electron acceptor (Figs. 4d,e), demonstrating the key role of MccSep in S^0^ respiration. Western blot analysis revealed a pronounced increase in MccSep protein levels upon S^0^ supplementation (Fig. 4f), consistent with our previous transcriptome data showing S^0^-induced transcriptional upregulation of *mccsep*^21^. These findings demonstrate the pivotal role of MccSep in S^0^ respiration in *G. sulfurreducens* PCA.

**Fig. 4.**
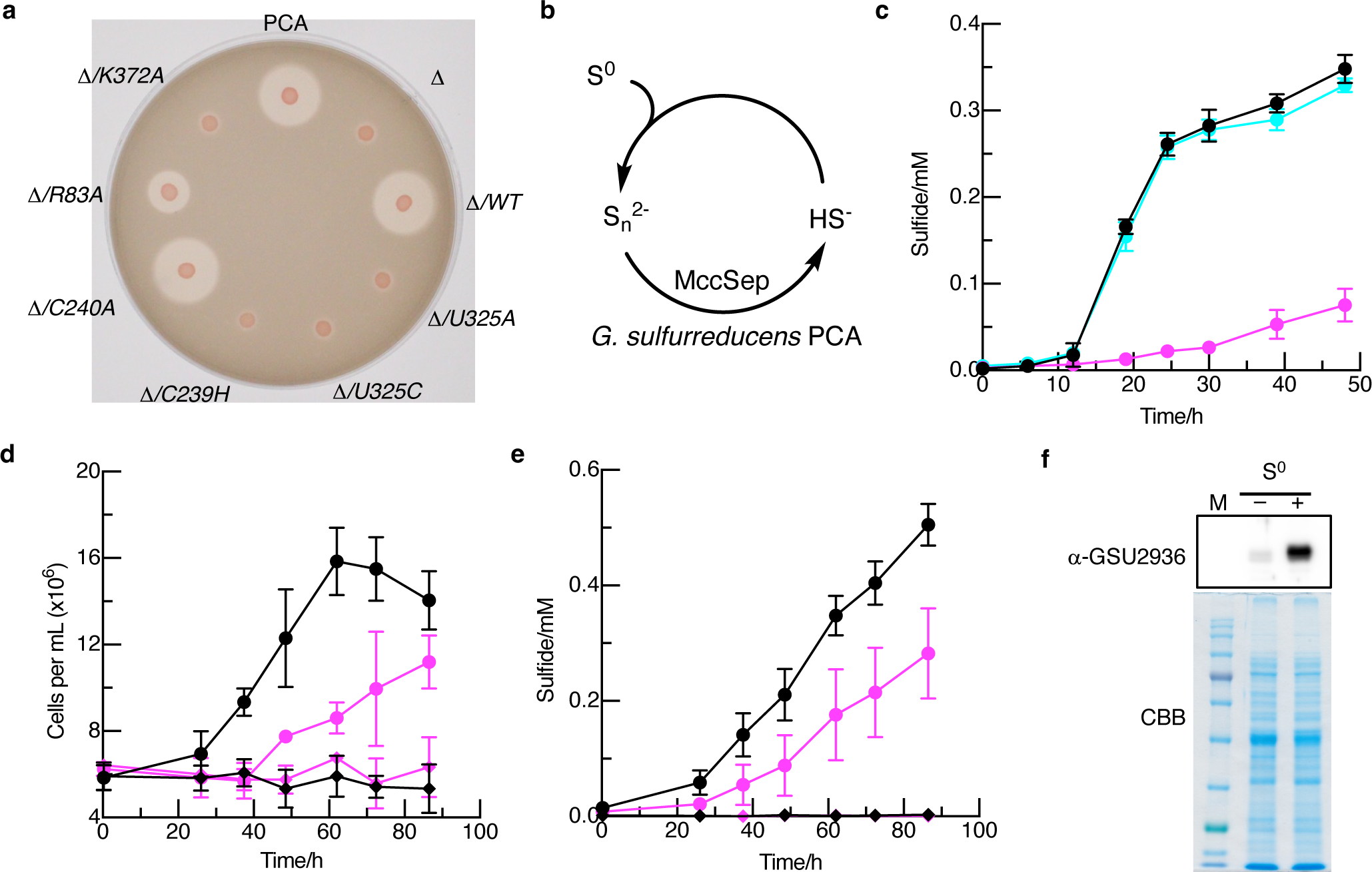
*In vivo* analyses of MccSep. **a**, Sulfur-reduction halo assay. The parent strain *G. sulfurreducens* PCA (PCA), the Δ*mccsep* strain (Δ), and Δ*mccsep* complemented with a plasmid expressing either the wild-type *mccsep* (WT) or its mutant (U325A, U325C, C239H, C240A, R83A, or K372A), were inoculated onto NBAFYE agar medium supplemented with S^0^ and grown anaerobically for 4 days. **b**, Schematic illustration of S^0^ reduction by *G. sulfurreducens* PCA, highlighting the essential role of MccSep in polysulfide (S_n_^2-^) reduction and HS^-^formation to facilitate S^0^ reduction. **c**, Sulfide production by the parent strain *G. sulfurreducens* PCA (black), Δ*mccsep* (magenta), and Δ*mccsep*/pMcasMccSep (cyan) cultivated in a liquid NBAFYE medium supplemented with 0.048% sulfur powder. Data presented are mean ± SD of triplicate experiments. **d**,**e**, Growth (**d**) and sulfide production (**e**) of the parent strain *G. sulfurreducens* PCA (black) and the Δ*mccsep* mutant (magenta) in the presence (circles) or absence (diamonds) of 0.1% (w/v) sulfur powder as a sole electron acceptor in a minimum liquid FWA medium containing 20 mM acetate as the electron donor. Data presented are mean ± SD of triplicate experiments. **f**, Induction of MccSep expression by S^0^ was examined by western blot analysis of crude extracts of the parent strain colonies formed on NBAFYE agar medium supplemented without (–) or with (+) S^0^. MccSep was detected using an anti-GSU2936 antibody (upper panel). The lower panel shows the identical gels stained with CBB to visualize the amount of proteins loaded.

## Discussion

Reductive cleavage of S–S bond is a central reaction in biology. The S–S bond cleavage of thioredoxin, glutathione, lipoamide, and coenzyme A is catalyzed by the flavoprotein disulfide reductase family, which employs at least one nonflavin redox center to transfer electrons from reduced pyridine nucleotide (NADPH/NADH) to substrate via a FAD cofactor^37^. Over the past three decades, enzymological studies on bacterial and archaeal S^0^/polysulfide reduction have revealed diverse strategies for S–S bond cleavage^2,38^. Molybdenum cofactor-dependent enzymes such as Psr^39^ and Sre^40^, the best-characterized bacterial sulfur reductases, are coupled with membrane-bound hydrogenases to use H_2_ as the electron donor. Cytosolic NAD(P)/FAD/CoA-dependent Npsr and NSR catalyze polysulfide or disulfide reduction, although their physiological roles remain unclear^41,42^. The archaeal sulfur respiratory complex MBS contains only iron-sulfur clusters^43^. Although the triheme cytochrome *c*_3_ (also known as *c*_551.5_ and *c*_7_) exhibits polysulfide-reducing activity^10^, its physiological relevance to sulfur respiration in S^0^-reducing bacteria has not been established. The discovery of the polysulfide reductase MccSep expands the chemical diversity of S–S bond-cleaving enzymes, defining a previously unrecognized class of redox catalysts that uniquely integrates a His/Cys-ligated heme with a Sec residue within the same catalytic framework. Sec plays a crucial role in various redox enzymes because the selenol group has a low p*K*_a_ (5.2), in contrast to the higher p*K*_a_ (8.5) of the thiol group in Cys^44^. The low p*K*_a_ of the selenol group allows it to remain deprotonated mainly at physiological pH, enhancing its nucleophilicity in redox reactions^44^. These properties render Sec exceptionally effective for catalysis^45^.

In MCC enzymes, the unoccupied coordination position at the active-site heme typically serves as a direct binding site for small substrate molecules such as nitrite, sulfite, and hydroxylamine^46^. Electrons are subsequently transferred through a closely linked chain of heme groups^14,47^. However, this reaction mechanism makes it challenging to discriminate among small inorganic substrates. Indeed, most MCC enzymes exhibit promiscuity toward nitrogen and sulfur substrates, as exemplified by the nitrite reductase NrfA, which acts on nitrite, hydroxylamine, and sulfite^47^, and the tetrathionate reductase OTR, which utilizes both nitrite and thiosulfate^48^. In MccSep, the corresponding coordination position of the active-site heme is occupied by the S_γ_ atom of C239 (Fig. 3c). We propose that this configuration may prevent non-specific binding of small inorganic anions to heme 2.

Based on the structural and biochemical evidence, we propose a catalytic mechanism for MccSep (Fig. 5). Upon substrate binding, U325 adopts the conf2 conformation, positioning its selenol group near the substrate for nucleophilic attack (Fig. 3c). U325 then shifts to conf1, bringing the Se_γ_ atom covalently attached to the substrate moiety closer to the S_γ_ atom of C239, which subsequently undergoes a nucleophilic attack to form the U325-Se_γ_–Se/S–S_γ_-C239 intermediate. The relatively long distances between the S_γ_ atom of C239 and the Se_γ_ atom of U325 in both conformations (conf1: 3.6 Å; conf2: 3.6 Å), together with the absence of continuous electron density between them, support this interpretation (Fig. 3c and Extended Data Fig. 6d). Finally, electron transfer from heme 2 reduces the U325-Se_γ_–Se/S–S_γ_-C239 intermediate, cleaving the bonds and completing the catalytic cycle. The dynamic conformational changes of U325 likely facilitate both the nucleophilic attack on the substrate and the subsequent positioning and reduction of the intermediate near the heme-ligating C239. Collectively, these features define a catalytic mechanism that shares conceptual similarities with mammalian selenoprotein thioredoxin reductase^49^, yet represents a distinct Sec- and heme-dependent mode of S–S bond cleavage. In addition, the mechanism of action of MccSep may be related to the reverse reaction of thiosulfate oxidation catalyzed by the MCC enzymes TsdA^16^ and SoxAX^50^, both of which employ His/Cys-liganded heme for catalysis.

**Fig. 5.**
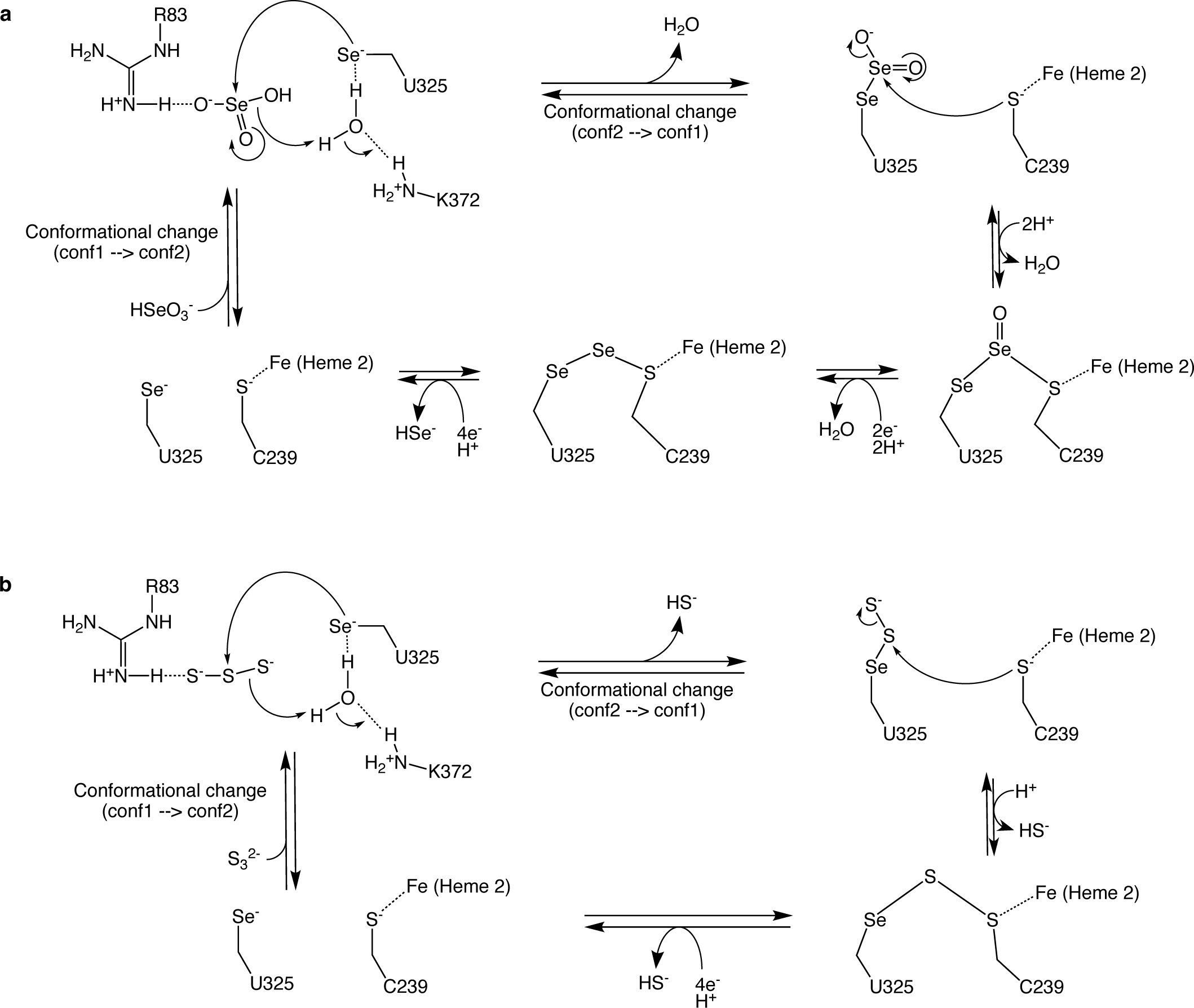
Proposed reaction mechanism for MccSep. **a**, Reaction mechanism for selenite reduction. **b**, Reaction mechanism for polysulfide reduction (illustrated using a representative trisulfide substrate).

While our data indicate that MccSep catalyzes polysulfide reduction in the periplasm, the route through which extracellularly generated polysulfides enter the periplasmic space remains unclear. The present findings suggest that the sulfide produced by MccSep may react with extracellular S^0^ to form polysulfides, which subsequently serve as substrates for MccSep (Fig. 4b). Although the transport mechanism for these sulfur species across the outer membrane has not been elucidated, the involvement of additional periplasmic or membrane-associated sulfur-transfer components cannot be ruled out. Elucidating how these sulfur intermediates are delivered to MccSep is important for future study.

## Methods

### Bacterial strains and growth conditions

*G. sulfurreducens* PCA^T^ (=DSM 12127^T^) was obtained from Leibniz Institute DSMZ-German Collection of Microorganisms and Cell Cultures (Braunschweig, Germany). Manipulations of the bacterium were performed inside an anaerobic chamber (Glovebox Japan Inc., Tokyo, Japan) under a nitrogen gas atmosphere. The *G. sulfurreducens* PCA cells were cultured in NBAFYE^51^ medium containing 20 mM acetate, 40 mM fumarate, 0.1% yeast extract, and 1 mM cysteine under strict anaerobic conditions at 35 °C. An S^0^-supplemented NBAFYE liquid medium was prepared by adding 0.048% (w/v) sulfur powder (Nacalai Tesque, Kyoto, Japan) to the NBAFYE medium before autoclaving. A minimum liquid medium supplemented with S^0^ was a modification of FWA-Fe(III) citrate medium^52^, containing 20 mM acetate as the electron donor and 0.1% (w/v) sublimed sulfur powder (Strem Chemicals, Newburyport, MA, USA) in place of iron(III) citrate, as the electron acceptor. To prepare the dispersed S⁰/polysulfide agar medium, a 2 mM polysulfide solution (prepared as described elsewhere) was added to NBAFYE medium containing 1% agar after autoclaving, and the medium was allowed to air-oxidize, forming dispersed S^0^ and releasing excess H_2_S after adjusting its pH to 7 with HCl. This procedure yields a suspension containing finely dispersed S⁰ and residual polysulfide species in equilibrium^33,34^. The dispersed S⁰/polysulfide NBAFYE plates were kept in an anaerobic gas-producing pouch (AnaeroPak Kenki, Mitsubishi Gas Chemical, Tokyo, Japan) for at least 16 h to remove oxygen before use. For the S^0^-reducing halo assay, bacterial cells (1 μL of OD_600_ = 1.0) were spotted on S^0^-supplemented NBAFYE plates supplemented with 1 mM isopropyl-β-D-thiogalactopyranoside and then incubated at 35 °C for 2–6 days. *E. coli* DH5α was used for general DNA manipulations, and *E. coli* BL21(DE3) (Novagen, Madison, WI, USA) was used to prepare the GSU2936 antigen. *E. coli* cells were aerobically grown in LB medium at 37 °C.

### Preparation of a polysulfide solution

A polysulfide solution was prepared by adding tetrathionate to an anaerobic solution of sulfide^33^. Briefly, 0.25 M Na_2_S_4_O_6_ in 0.5 M Tris-HCl (pH 8.5) was anaerobically added to 1 M Na_2_S in 0.5 M Tris-HCl (pH 8.5) at a 1:1 ratio. The resulting solution contained a mixture of polysulfide species of various chain lengths (S_n_^2-^), and its concentration was determined at 360 nm (ε = 0.38 cm⁻¹ mM⁻¹ per sulfur atom)^33^ and expressed as total sulfur equivalents.

### Preparation of an anti-GSU2936 antibody

To obtain the GSU2936 protein as an antigen, a DNA fragment containing the *gsu2936* gene was cloned using primers GSU2936-F and GSU2936-R (Supplementary Table 1) and inserted into the NdeI and XhoI sites of pET21a(+) (Takara Bio Inc., Kusatsu, Shiga, Japan) to yield pETGSU2936. *E. coli* BL21(DE3) cells carrying pETGSU2936 were cultured in LB with 100 μg/mL of ampicillin at 25 °C for 5 h. After induction with 1 mM isopropyl-β-D-thiogalactopyranoside for 6 h, the cells were harvested by centrifugation at 5,000 × *g*. Under the conditions, a GSU2936 peptide was expressed as an inclusion body. After sonication and centrifugation, the pellet was treated as previously described^53^. GSU2936 was isolated from an SDS-PAGE gel and used to immunize a rabbit. Antisera obtained from the immunized rabbit were used as an anti-GSU2936 antibody.

### Purification of native MccSep from *G. sulfurreducens* PCA

*G. sulfurreducens* PCA was cultured in NBAFYE medium containing 17 mM sodium fumarate and 0.055% (w/v) sulfur powder (Nacalai Tesque) at 35 °C for 40 h. Cells were harvested by centrifugation, suspended in buffer A (20 mM Tris-HCl, 1 mM DTT, pH 7.8), and disrupted by sonication. Ammonium sulfate (final conc. 1.75 M) was added to the crude extract, kept at 4 °C for 3 h, and centrifuged. The supernatant was filtered and applied to a Butyl-Toyopearl (Tosoh Corp., Tokyo, Japan) column equilibrated with buffer A containing 1.75 M ammonium sulfate. After washing with buffer A containing 1.5 M ammonium sulfate, the protein was eluted with a linear gradient of 1.5-1.0 M ammonium sulfate in buffer A. MccSep-containing fractions, identified by western blot using anti-GSU2936 antibody, were collected. After ammonium sulfate was added to the collected solution at 1.4 M, the protein was applied to a second Butyl-Toyopearl column equilibrated with buffer A containing 1.4 M ammonium sulfate. Proteins were eluted with buffer A containing 1.2 M ammonium sulfate. MccSep-containing fractions were pooled, concentrated, and dialyzed against buffer A. The protein solution was applied into a Q-Sepharose column (Cytiva, Marlborough, MA, USA) equilibrated with buffer A. After washing with buffer A containing 170 mM NaCl, the proteins were eluted with buffer A containing 250 mM NaCl. The MccSep-containing fractions were collected and dialyzed against buffer A.

### Trypsin digestion, HPLC, and ESI–MS analysis

Purified MccSep (0.25 mg/mL) was denatured and reduced in 0.22 mL of 0.27 M Tris-HCl (pH 8.5) containing 6 M guanidine hydrochloride and 4 mM dithiothreitol. The protein was carboxymethylated with 8 mM iodoacetamide in the dark for 1 h. After adding 1.2 mL of ice-cold methanol/chloroform/water (4:1:3), the solution was centrifuged to form a protein aggregate at the interface. The upper layer was removed, 0.45 mL of cold methanol was added, and the mixture was centrifuged. The protein pellet was lyophilized, dissolved in 80 mM Tris-HCl buffer (pH 8.5) containing 1.2 M urea, and digested with trypsin (2.3 μg/mL) at 37 °C for 16 h. The reaction was stopped with formic acid and centrifuged, and the supernatant was applied to reversed-phase HPLC using a Capcell Pak C_18_ SG120 column (Shimadzu, Kyoto, Japan) and a Shimadzu HPLC system. The peptides were eluted with an increased acetonitrile concentration in a solvent containing 0.075% formic acid. The peptide fractions were analyzed by Edman sequence analysis with Shimadzu PPSQ-21A protein sequencer (Shimadzu) and ESI–MS with a triple-quadrupole Sciex API 3000 mass spectrometer (Applied Biosystems, Foster City, CA, USA) equipped with an electrospray ionization source in positive mode. ESI–MS analysis indicated that peptides eluted at 13-14 min included the target Sec-containing peptide (VEMTNHAGHSIPDG<u>U</u>PTPNR) with *Se*-carbamidomethylation on Sec (calculated mass of 2237.8 Da). This fraction was further purified by a second HPLC (Shimadzu). A well-separated peptide eluted at 22.3 min was analyzed by ESI–MS (Applied Biosystems).

### Determination of selenium, iron, sulfide, and protein

Selenium content in purified MccSep was determined fluorometrically after acid digestion and derivatization using 2,3-diaminonaphthalene^27^. Iron content in MccSep was determined colorimetrically using the iron-bathophenanthroline disulfonate complex formation^26^. Sulfide in a culture solution was determined by a methylene blue method as previously described^54^. Protein concentrations were measured using a Protein Assay BCA kit (Nacalai Tesque).

### Subcellular fractionation, SDS-PAGE, and western blot analyses

Subcellular fractionation was performed as previously described^55^. SDS-PAGE analyses used 10–12% polyacrylamide gels, with proteins stained with Coomassie Brilliant Blue G-250. Western blotting employed Immobilon-P polyvinylidene difluoride membranes (0.45-μm pore size; Merck Millipore, Burlington, MA, USA) and a Trans-Blot SD semi-dry transfer cell (Bio-Rad, Hercules, CA, USA). Immunoreactive proteins were detected using anti-rabbit IgG(H+L) HRP-conjugated secondary antibody (Promega, Madison, WI, USA), Chemi-Lumi One Super (Nacalai Tesque), and an Amersham Imager 600 (Cytiva). Molecular mass size markers included Protein Markers for SDS-PAGE (Nacalai Tesque), BLUeye Prestained Protein Ladder (Sigma-Aldrich, St. Louis, MO, USA), MagicMark XP Western Protein Standard (Thermo Fisher Scientific, Waltham, MA, USA), and WIDE-VIEW Prestained Protein Size Marker III (Fujifilm Wako Pure Chemical Co., Osaka, Japan).

### Construction of Δ*mccsep*

Primers used are listed in Supplementary Table 1. The *mccsep*-deficient mutant with a kanamycin-resistant gene cassette (*kanR*) insertion was constructed by homologous gene replacement (Extended Data Fig. 2a). A DNA region spanning 276 bp upstream to 231 bp downstream of the ATG initiation codon of *mccsep* was PCR-amplified from *G. sulfurreducens* PCA genomic DNA using primers GSU01 and GSU06. A DNA region ranging from 118 bp upstream to 450 bp downstream of the TAG stop codon of *mccsep* was amplified using primers GSU05 and GSU02. A 1149 bp *kanR* fragment was amplified from pJRD215^56^ using primers GSU03 and GSU04, which have sequences complementary to GSU06 and GSU05, respectively. These three fragments were purified and fused by PCR using primers GSU01 and GSU02. The resulting DNA fragment was introduced into *G. sulfurreducens* PCA by electroporation (GTE-10, Shimadzu), and disruptants were selected on NBAFYE agar medium with 400 μg/mL kanamycin. Correct gene disruption was confirmed by PCR (Extended Data Figs. 2b,c), and the absence of the MccSep protein in Δ*mccsep* was confirmed by western blot analysis of crude extracts (Extended Data Fig. 3a). RT-PCR analysis of *gsu2935*, located downstream of *mccsep*, suggested no negative polar effect on downstream genes (Extended Data Fig. 2d and Supplementary Method 1).

### Construction of expression plasmids for WT and mutants of MccSep

Primers used are listed in Supplementary Table 1. To express MccSep with a C-terminal Strep-tag in Δ*mccsep*, a DNA cassette containing a ribosome-binding site and restriction sites (NdeI, XhoI, and XbaI) was prepared using oligonucleotides casF and casR and inserted into the EcoRI site of pMMB206^57^ to yield pMcas. A DNA fragment containing *mccsep* was amplified from genomic DNA by PCR using primers MccSep-F and MccSep-Stag-R to introduce sequences for Factor Xa cleavage and Strep-tag sites. This fragment was re-amplified using the primers MccSep-F and Stag-Xba-R to introduce an XbaI site and inserted into pMcas to yield pMcasMccSepStag. To express MccSep without a tag for genetic complementation, a *mccsep* fragment was amplified from genomic DNA using primers MccSep-F and MccSep-R and inserted into pMcas to yield pMcasMccSep. To produce U325A, U325C, and C240A mutants, the QuikChange site-directed mutagenesis method (Agilent Technologies, Santa Clara, CA, USA) was used with appropriate primers. The sequences of constructed plasmids were verified by DNA sequencing (ABI Genetic Analyzer 3130, Thermo Fisher Scientific). Plasmids expressing each of R83A, C239H, and K372A were constructed based on pMcasMccSepStag by GenScript. Plasmids were introduced into bacterial cells by electroporation using either GTE–10 (Shimadzu) or NEPA Porator (Nepa Gene, Chiba, Japan).

### Purification of recombinant enzymes

The Δ*mccsep* cells harboring pMcasMccSepStag were anaerobically grown in NBAFYE medium containing 10 μg/mL chloramphenicol at 35 °C for 24 h. After induction of gene expression with 1 mM isopropyl-β-D-thiogalactopyranoside, the cells were grown for 24 h at 30 °C and harvested by centrifugation. The cells were suspended in 0.1 M Tris-HCl buffer (pH 8.0) containing 0.3 M NaCl, 5 mM 2-mercaptoethanol, and 0.1% Triton X-100 and disrupted by sonication. The cell debris was removed by centrifugation, and the supernatant solution was applied to a Strep-Tactin Superflow column (IBA GmbH, Göttingen, Germany). The enzyme was eluted with varied concentrations of D-desthiobiotin. The enzyme fractions were concentrated with an Amicon Ultra 30K (Merck Millipore) and applied to a Superdex 200 Increase 10/300 GL column (Cytiva) equilibrated with 30 mM Tris-HCl buffer (pH 7.5) containing 500 mM NaCl, using an ÄKTA go chromatography system (Cytiva) at 0.5 mL/min. The enzyme fractions were collected, concentrated, and buffer-exchanged to 10 mM HEPES-NaOH (pH 7.5) with an Amicon Ultra 30K. For crystallization, further polishing was carried out with a Superdex 200 Increase 10/300 GL column equilibrated in 10 mM HEPES-NaOH buffer (pH 7.5). The purified enzyme was stored at –80 °C until use. Mutant enzymes were purified similarly.

### Crystallization and data collection

Purified WT and U325A mutant enzymes were concentrated to 5–10 mg/mL in 10 mM HEPES-NaOH (pH 7.5). Initial screening of crystallization conditions was performed by the sitting-drop vapor-diffusion method using a crystallization robot, mosquito (SPT Labtech, Melbourn, UK). Several crystals were obtained from commercially available screening kits, JCSG Plus, PACT Premier, NeXtal Classics suite, ProPlex (Molecular Dimensions, Sheffield, UK), and MembFac (Hampton Research, Aliso Viejo, CA, USA). After optimization of the crystallization reservoir solution, single crystals of both WT and U325A mutant suitable for X-ray data collection were obtained by hanging-drop vapor-diffusion method using a reservoir solution containing 0.1 M ammonium sulfate, 0.1 M HEPES-NaOH (pH 7.5), and 10% PEG4000. Iodine-derived crystals of WT MccSep were prepared by the vaporizing iodine labeling method^58^. A droplet of iodine solution (0.67 M KI and 4.7 M I_2_) was placed next to the crystallization droplet containing MccSep crystals. After 20 min, the iodine solution was removed and the crystallization droplet was incubated for a further 20 h. Prior to the X-ray experiment, the crystal was cryo-protected by the addition of 20% ethylene glycol and flash-cooled by an N_2_-gas stream at 100 K. X-ray diffraction data sets were collected at the SPring-8 beamline 26B1 (Hyogo, Japan) equipped with an Eiger 4M detector (Dectris, Baden-Dättwil, Switzerland) at wavelengths 0.9790 Å (native) and 1.9000 Å (iodine derivative).

### Structure determination and refinement

Diffraction data sets were indexed, integrated, and scaled using XDS ver. Jan 26, 2018^59^. The crystal structure of WT MccSep was solved by the single isomorphous replacement with anomalous scattering (SIRAS) method using native and iodine-derivative data sets with the program Autosol in PHENIX ver. 1.20.1_4487^60^. Phase calculation resulted in good-quality electron density maps as judged by the figure of merit (0.45) and the successful autotracing of the four MccSep copies in the asymmetric unit. Several rounds of refinement using phenix.refine and manual model building with Coot ver. 0.8.9^61^ were performed. The structure of the U325A MccSep mutant was phased by molecular replacement (MR) using the wild-type MccSep model with the program phenix.MRage in PHENIX ver. 1.20.1_4487^60^. Stereochemical parameters were checked using Molprobity ver. 4.5.2^62^ in PHENIX ver. 1.20.1_4487^60^. Ramachandran statistics (percentages of favored/allowed/disallowed) are 96.82/3.18/0.00 for WT, 96.94/3.00/0.06 for the U325A, 96.40/3.60/0.00 for the WT-selenite complex, 96.40/3.60/0.00 for the WT-selenate complex, and 95.68/4.32/0.00 for the U325A-selenate complex. The figures were prepared using PyMOL ver. 2.4.0 (Schrödinger, LLC, USA).

### Enzyme assays and kinetics

The activity of MccSep with polysulfide and selenite was determined in triplicate under anaerobic conditions using 1.0 mL quartz cuvettes sealed with gas-tight rubber stoppers. Reactions were monitored at 600 nm with a Shimadzu UV-1850 UV–vis spectrophotometer at 30 °C, based on the linear decrease in absorbance due to the oxidation of reduced MV (MV^•+^). Unless otherwise noted, each reaction mixture contained 0.3 mM methyl viologen, 0.075 mM dithionite, 2 mM CaCl_2_, 0.2 mM substrate, and 0.8–5 μg/mL of MccSep in 50 mM Bicine-NaOH buffer (pH 8.5). For pH-dependent activity assay with selenite, the reaction conditions were modified to include 0.8 mM MV^•+^, 0.17 mM dithionite, and 10 mM selenite. The enzymatic reaction rates were corrected by subtracting background values obtained from control reactions lacking the enzyme. Activity was calculated using the extinction coefficient of MV^•+^ (*χ*_600_ = 13.7 mM^-1^ cm^-1^)^63^ and expressed as μmoles of MV^•+^ oxidized per min per mg of protein. Kinetic parameters were determined by varying the substrate concentration under the conditions described above. Nonlinear curve fitting of the experimental data was performed using GraphPad Prism ver. 10.6.1 for MacOS (GraphPad Software, San Diego, CA, USA), and kinetic parameters were derived based on an allosteric-sigmoidal model: *v* = *V*_max_[S]*^h^*/(*K*_0.5_*^h^* + [S]*^h^*), where *v* is the reaction velocity, *V*_max_ is the maximum velocity, [S] is the substrate concentration, *h* is the Hill coefficient, and *K*_0.5_ is the substrate concentration at a half-maximal velocity^64^.

### Bioinformatics analysis

The protein sequence retrieval for MccSep homologs was performed using the DIAMOND BLASTp search ver. 2.1.10^65^ with an ultra-sensitive mode from the National Center for Biotechnology Information (NCBI) non-redundant protein sequence database (November, 2024)^66^ using *G. sulfurreducens* MccSep as a query sequence. Short-length (amino acid length <200) and low-score (e-value >0.01) hits were excluded, and then the sequences with deletions or mutations in the conserved heme-binding motifs and fragments that were truncated before −1 position from the Sec site were filtered out by verifying the multiple sequence alignment prepared using the MAFFT ver. 7.487 program with the E-INS-i method^67^. In addition, the MccSep homologs-encoding genomes were retrieved from the NCBI assembly database (December, 2024) by searching for protein accession numbers. The protein sequences without genomic information were excluded. For the protein sequences stopped at -1 position from the Sec site in the multiple sequence alignment, 900 bp downstream sequence of the TGA stop codon of the corresponding nucleic acid sequence was extracted using SeqKit ver. 2.9.0^68^, and then the open reading frame of the C-terminal fragment was translated using the getorf command in the EMBOSS package ver. 6.6.0^69^. Finally, a Sec-containing full-length sequence was obtained by combining the TGA-stopped N-terminal fragment with Sec inserted to the N-terminus. After sequence clustering of all identified 637 non-redundant MccSep homologs containing the conserved Sec/Cys residue at 90% sequence identity using the UCLUST algorithm in USEARCH ver. 11.0.667^70^, the 455 centroid sequences of the clusters were aligned using the MAFFT program with the E-INS-i method. Ambiguously aligned sites were subsequently trimmed using the trimAl ver. 1.4.1 program with the “automated1” mode^71^. The trimmed alignment for phylogenetic analysis contained 283 amino acid sites that were used for the maximum likelihood phylogenetic tree reconstruction using IQ-TREE ver. 2.3.6^72^ with the Q.pfam+R8 model assigned by ModelFinder^73^. The reliability of the tree topology was evaluated by the SH-aLRT^74^ and UFBoot support values based on 1,000 resamplings^75^. The trees were visualized using iTOL ver. 7.2^76^. Genome-based taxonomies of the 684 MccSep homologs-encoding genomes were annotated using GTDBTk ver. 2.4.0 with the GTDB database release R220^77^. RNA secondary structure was predicted by the MXFold2 server^78^ using the SECIS region consisting of the UGA and its 30 nt downstream sequence. The subcellular localization was predicted by DeepLocPro ver. 1.0^25^.

## Data availability

All atomic coordinates for MccSep proteins have been deposited in the Protein Data Bank (PDB) under accession codes 9WT8, 9WT9, 9WTA, 9WTB, and 9WTC.

## Acknowledgments

We are grateful to Ms. Erina Shinno (Ritsumeikan University), Ms. Miki Jinno (Ritsumeikan University), Mr. Bunta Yoshimura (Ritsumeikan University), Dr. Ishrat Mst. Jahan (Ritsumeikan University), Dr. Yasushi Tani (Ritsumeikan University), Dr. Anna Ochi (Ritsumeikan University), Dr. Shigeki Saito (Ritsumeikan University), and Dr. Jun Kawamoto (Kyoto University) for valuable technical assistance, and Professor Nobuyoshi Esaki (Kyoto University) for providing expert advice from a broad scientific perspective.

## Author information

Takuya Yoshizawa

Present address: Research Division, Chugai Pharmaceutical Co., Ltd, Yokohama, Kanagawa, Japan

Ryuta Tobe

Present address: Graduate School of Agricultural Science, Tohoku University, Sendai, Miyagi, Japan

Authors and Affiliations

**College of Life Sciences, Ritsumeikan University, Kusatsu, Shiga, Japan**

Hisaaki Mihara, Takuya Yoshizawa, Yukiko Izu, Nana Shimamoto, Masao Inoue, Ryuta Tobe, Riku Aono, and Hiroyoshi Matsumura

**Ritsumeikan Global Innovation Research Organization, Ritsumeikan University, Kusatsu, Shiga, Japan**

Masao Inoue

**Institute for Chemical Research, Kyoto University, Uji, Kyoto, Japan**

Wanjiao Zhang & Tatsuo Kurihara

## Contributions

H.M.(Mihara), W.Z., T.K., and T.Y. designed the research. W.Z. performed the purification and biochemical analyses of the native enzyme. W.Z., N.S., and R.T. constructed the plasmids. N.S. and R.T. performed the gene disruption and RT-PCR experiments. Y.I. performed most experiments using purified recombinant enzymes and gene-disruptant strains. T.Y. and H.M.(Matsumura) performed the crystallographic experiments. M.I. performed bioinformatic analysis. H.M.(Mihara), T.Y., Y.I., W.Z., M.I., T.K., R.T., R.A., and H.M.(Matsumura) analyzed data. H.M.(Mihara), H.M.(Matsumura), R.T., and M.I. wrote the paper with comments from R.A.

## Corresponding authors

Correspondence to Hisaaki Mihara or Hiroyoshi Matsumura.

## Extended data figures and tables

**Extended Data Fig. 1 Nucleotide and amino acid sequences of MccSep.**

The 81-nt region flanking *gsu2937* and *gsu2936* is underlined. The TGA codon encoding selenocysteine (U) at the amino acid position 325 is shown in blue. The erroneously annotated start codon of *gsu2936* and the corresponding methionine residue are indicated by magenta. The postulated SECIS element is indicated by underlined red letters. The N-terminal signal peptide (1–25) is shaded in gray. Five heme-binding motifs (CxxCH) and heme numbering are highlighted in cyan. The nucleotide and amino acid sequences of MccSep were retrieved from the NCBI database (accession numbers: GS_RS14775 and WP_262421250.1, respectively).

**Extended Data Fig. 2 Construction of an *mccsep*-deficient strain (Δ*mccsep*).**

**a**, Schematic drawing of the homologous recombination between the disruption construct and the genomic DNA of the parent strain *G. sulfurreducens* PCA (PCA). Primer annealing sites are indicated. **b**, Schematic drawing of the corresponding genomic region of Δ*mccsep.* Primer annealing sites and predicted sizes of PCR fragments are indicated. **c**, PCR analysis using genomic DNAs from the parent PCA strain (lanes 1, 3, and 5) and Δ*mccsep* (lanes 2, 4, and 6). Primers used were GSU3219F and GSU5417R (lanes 1 and 2), GSU03 and GSU04 (lanes 3 and 4), and GSU01 and GSU04 (lanes 5 and 6). Asterisks indicate non-specific bands. **d**, Reverse transcription-PCR analysis showing no negative polar effect on the downstream gene expression by the *mccsep* disruption. mRNA levels of *gsu2935* and 16S rRNA in the parent PCA and Δ*mccsep* strains were analyzed by RT-PCR using each primer set (gsu2935-F/gsu2935-R and 16SrRNA-F/16SrRNA-R, respectively) shown in Supplementary Table 1. Cells were grown in NBAFYE medium at 35 °C to the mid-exponential phase.

**Extended Data Fig. 3 Expression and purification of recombinant MccSep.**

**a**, Western blot analysis of the crude extracts from the parent strain *G. sulfurrreducens* PCA (lane 1), Δ*mccsep* (lane 2), and Δ*mccsep* complemented with pMcasMccSepStag carrying the *mccsep* gene (lane 3). Crude extracts of *G. sulfurreducens* strains grown in NBAYFE medium were separated on 10% SDS-PAGE gels and analyzed by western blotting using anti-GSU2936 antibody (upper panel). The lower panel represents the identical gel visualized by CBB staining to show the amount of proteins loaded. **b**, Purification of recombinant MccSep (lane 1) and its mutants (lanes 2, U325A; 3, U325C; 4, C239H; 5, C240A; 6, R83A; and 7, K372A). Each recombinant enzyme was purified as a Strep-tag fusion protein and analyzed by SDS-PAGE. Proteins were visualized by CBB staining. **c**, UV–vis absorption spectra of air-oxidized (blue line) and 0.02 mM sodium dithionite-reduced (orange line) MccSep (0.06 mg/mL).

**Extended Data Fig. 4 Comparison with multiheme-binding proteins.**

Structural comparison of MccSep with representative multiheme-binding cytochromes. Upper panels show monomeric structures; lower panels display superimposed heme groups aligned with those of MccSep. **a**, MccSep. **b**, Penta-heme cytochrome *c*_552_ from *Thermochromatium tepidum* (PDB ID: ZE85). **c**, Octaheme sulfite reductase MccA from *Wolinella succinogenes* (PDB ID: 4RKM).

**Extended Data Fig. 5 Heme binding sites in MccSep.**

Hemes 1 (**a**), 3 (**b**), 4 (**c**), and 5 (**d**) are coordinated by two histidine residues, forming a bis-histidine axial ligation.

**Extended Data** Fig. 6 **Assignment of alternative conformations of the 323–326 loop.**

**a**, Initial refinement of the WT MccSep structure showing ambiguous electron density near U325. Negative *F_o_* – *F_c_* electron density peaks are shown on the main chain of G324 (red arrowhead) and the side chain of U325 (blue) with a positive peak *∼*3.0 Å from the oxygen atom of a sulfate ion (green). **b**, Initial refinement of the U325A mutant showing that the loop adopts a single conformation with clear electron density, except for a residual positive peak near C239. **c**, Superposition of the WT and U325A structures revealing differences in the C_α_–C_β_ orientation of residue 325, the main chain of G324, and the carbonyl group of D323. **d**, Partially overlapping 2*F_o_* – *F_c_* electron densities in the WT MccSep structure with those in the U325A structure suggest alternative loop conformations in WT. Refinement of the WT structure with dual conformations improved map interpretability. All maps: 2*F_o_* – *F_c_* (blue, 1.2σ); *F_o_* – *F_c_* positive (green, 4σ); *F_o_* – *F_c_* negative (red, –3σ).

**Extended Data Fig. 7 Anomalous difference Fourier maps at the active site.**

**a–c**, Anomalous difference Fourier maps (magenta, 2.5σ, λ = 0.9792 Å) for WT MccSep soaked with 10 mM selenite overnight (**a**), 10 mM selenate overnight (**b**), and for U325A soaked with 100 mM selenate (**c**). Anomalous difference Fourier map for WT MccSep soaked with 10 mM selenite (**a**) shows a continuous electron density between U325 in conf2 and bound selenite.

**Extended Data Table 1 Kinetic parameters for polysulfide reduction by MccSep and its mutants**

**Extended Data Table 2 Data collection, phasing, and refinement statistics**

## Supplementary information

Supplementary Information

This file contains Supplementary Table 1 and Supplementary Method 1.

Supplementary Data

This file contains Supplementary Data 1–3.

Supplementary Dataset

This file contains Supplementary Datasets 1 and 2.

