## Extended Data Figures for "Multiheme selenoenzyme essential for elemental sulfur respiration"

```
   1 ATGAAGCACAAAACCGCATGGCTGACCCTTGCTGCCGCAGCGCTTGCGCTCTGTGCCGCCGCACCGGTTTTTGCCGAAAAGGCGGGCATCGGCTGGCAGGAGACG
   1  M  K  H  K  T  A  W  L  T  L  A  A  A  A  L  A  L  C  A  A  A  P  V  F  A  E  K  A  G  I  G  W  Q  E  T

 106 ATCGTTGCCAAGAGCGGCAAGGCAAAGACCATGGCCGAGTTGGCAAAGATGTATGATTCCAGCTCCTGCATCGAGTGTCACCAGGAAGTCCACGATGAATGGGAG
  36  I  V  A  K  S  G  K  A  K  T  M  A  E  L  A  K  M  Y  D  S  S  S  C  I  E  C  H  Q  E  V  H  D  E  W  E
                                                                     Heme 1
 211 CAGTCGATCCATGCCCGCTCAATCTTCGGTACCGGCCGGACTGCCGCCACCTTCATGACTGCTGTGGTCAACGGCCTCATGGAATGGGAATATTCGGGGGTGAAA
  71  Q  S  I  H  A  R  S  I  F  G  T  G  R  T  A  A  T  F  M  T  A  V  V  N  G  L  M  E  W  E  Y  S  G  V  K

 316 AGCCCGTCGGACGTGAAAGTTGAGCACCTGATGGGATGCGCCAAGTGCCATCTCCCCCAACTGGCCGACGCCGAGGACTCGGTGGCCAAGGAGATCATCTCCACC
 106  S  P  S  D  V  K  V  E  H  L  M  G  C  A  K  C  H  L  P  Q  L  A  D  A  E  D  S  V  A  K  E  I  I  S  T
                                          Heme 2
 421 ATCGGCAGCTGGCAGGATGCACTGAGGAAAAAGGATTCGGCAAAGGCGGTTGAAGAGGCCGACAAGCTCAAGAGCCTCAACATCAATTGCCTCGTTTGCCATAAC
 141  I  G  S  W  Q  D  A  L  R  K  K  D  S  A  K  A  V  E  E  A  D  K  L  K  S  L  N  I  N  C  L  V  C  H  N
                                                                                              Heme 3
 526 CGTAACGCCATCACCCACAAGTGGACCGACGGCTATCCCCAGGCCGGGGTAGTCTACGGCTCCAAGGAAGGGGAACACCCCTCTGCGGCATTCCCCACGATGAAG
 176  R  N  A  I  T  H  K  W  T  D  G  Y  P  Q  A  G  V  V  Y  G  S  K  E  G  E  H  P  S  A  A  F  P  T  M  K

 631 GTCAGTCCGATCATGTCAGAGTCCATCCAGTGCGGCCAATGCCACGGCTTGGGGCCGAACCTGGAATTGGATAACCCGACCCAGTGCTGCACCAGCTATGCGAGC
 211  V  S  P  I  M  S  E  S  I  Q  C  G  Q  C  H  G  L  G  P  N  L  E  L  D  N  P  T  Q  C  C  T  S  Y  A  S
                                    Heme 4
 736 TACCTCTGGGCCTACAAGGCCGAGGGCGGGCGGGAAAACTGCCAGGAATGCCACATGAAGAAGAGCAAGCTCGGGCACAACATGCAGAGCTACCGTGACCCGGGC
 246  Y  L  W  A  Y  K  A  E  G  G  R  E  N  C  Q  E  C  H  M  K  K  S  K  L  G  H  N  M  Q  S  Y  R  D  P  G
                                                Heme 5
 841 ATGGCCAAGGCCGCCGTTGAGTTCAAGGCCGAGGCCTATGGCTATCACTGGCGGGACGGGGCGCTGGTGACGCCCAAGGCAGTGGTGAAGGTCGAGATGACCAAT
 281  M  A  K  A  A  V  E  F  K  A  E  A  Y  G  Y  H  W  R  D  G  A  L  V  T  P  K  A  V  V  K  V  E  M  T  N

 946 CACGCGGGCCACTCGATCCCCGATGGCTGACCGACCCCCAACCGACTGGTTCTGTCGGTCATTGCAAAGACAAAAGACGGCGAAGAAGTGTTCAATCAGGAAAAG
 316  H  A  G  H  S  I  P  D  G  U  P  T  P  N  R  L  V  L  S  V  I  A  K  T  K  D  G  E  E  V  F  N  Q  E  K

1051 ATCTACATGCCGGTGCCCCAGCAGCTGGCCCGGGGTGACCGGATGGGGCGCGGCCCCTATGAGAAAAGCGGTATGATCGAGGATACGGGTCTGCCGCCGGGCAAG
 351  I  Y  M  P  V  P  Q  Q  L  A  R  G  D  R  M  G  R  G  P  Y  E  K  S  G  M  I  E  D  T  G  L  P  P  G  K

1156 AAGATCCACGAGCGCTTCGATATCCTCTTCCCCACCGAGGATGTGGTGGAGGACGGGAAGAAGGTTCGCAAGACCCTGGCCCACGATCTGGAAGTGGAGGTGAAA
 386  K  I  H  E  R  F  D  I  L  F  P  T  E  D  V  V  E  D  G  K  K  V  R  K  T  L  A  H  D  L  E  V  E  V  K

1261 CTCTGGTATCTCCCCTTCGGTTCAATGAATTCCGATCCGTTCCTCTGGCATGAGTTCACCCAGAAGGTGAGCATCAGCGCCAAGGGCAAGTAG
 421  L  W  Y  L  P  F  G  S  M  N  S  D  P  F  L  W  H  E  F  T  Q  K  V  S  I  S  A  K  G  K  *
```

**Extended Data Fig. 1 Nucleotide and amino acid sequences of MccSep.**
The 81-nt region flanking *gsu2937* and *gsu2936* is underlined. The TGA codon encoding selenocysteine (U) at amino acid position 325 is shown in blue. The erroneously annotated start codon of *gsu2936* and the corresponding methionine residue are indicated by magenta. The postulated SECIS element is indicated by underlined red letters. The N-terminal signal peptide (1–25) is shaded in gray. Five heme-binding motifs (CXXCH), along with heme numbering, are highlighted in cyan. The nucleotide and amino acid sequences of MccSep were retrieved from the NCBI database (accession numbers: GS_RS14775 and WP_262421250.1, respectively).

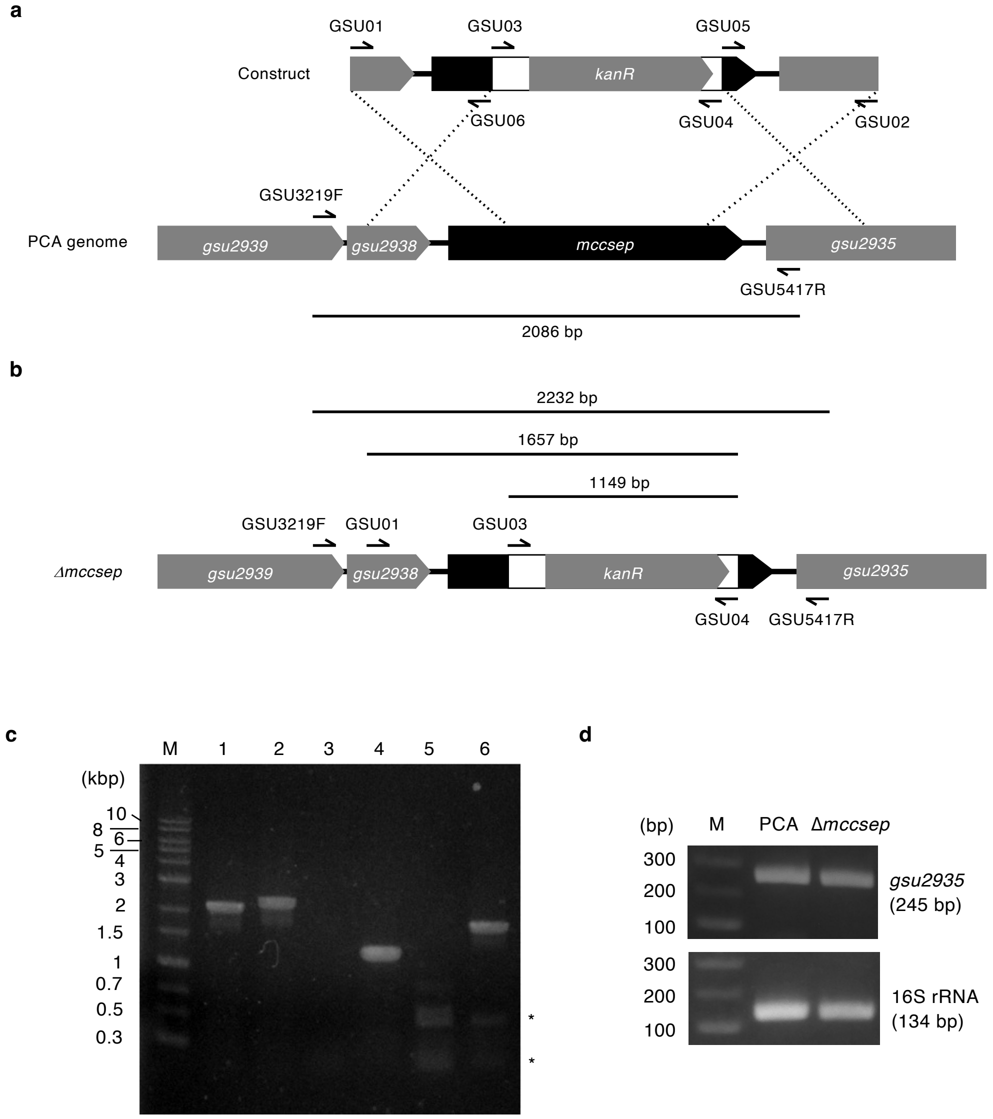


**Extended Data Fig. 2 Construction of an *mccsep*-deficient strain (Δ*mccsep*).**
**a,** Schematic drawing of the homologous recombination between the disruption construct and the genomic DNA of the parent strain *G. sulfurreducens* PCA (PCA). Primer annealing sites are indicated. **b,** Schematic drawing of the corresponding genomic region of Δ*mccsep.* Primer annealing sites and predicted sizes of PCR fragments are indicated. **c,** PCR analysis using genomic DNAs of the parent PCA strain (lanes 1, 3, and 5) and Δ*mccsep* (lanes 2, 4, and 6). Primers used were GSU3219F and GSU5417R (lanes 1 and 2), GSU03 and GSU04 (lanes 3 and 4), and GSU01 and GSU04 (lanes 5 and 6). The asterisks indicate non-specific bands. **d,** Reverse transcription-PCR analysis showing no negative polar effect on the downstream gene expression by the *mccsep* disruption. mRNA levels of *gsu2935* and 16S rRNA in the parent PCA and Δ*mccsep* strains were analyzed by RT-PCR using each primer set (gsu2935-F/gsu2935-R and 16SrRNA-F/16SrRNA-R, respectively) shown in Supplementary Table S1.

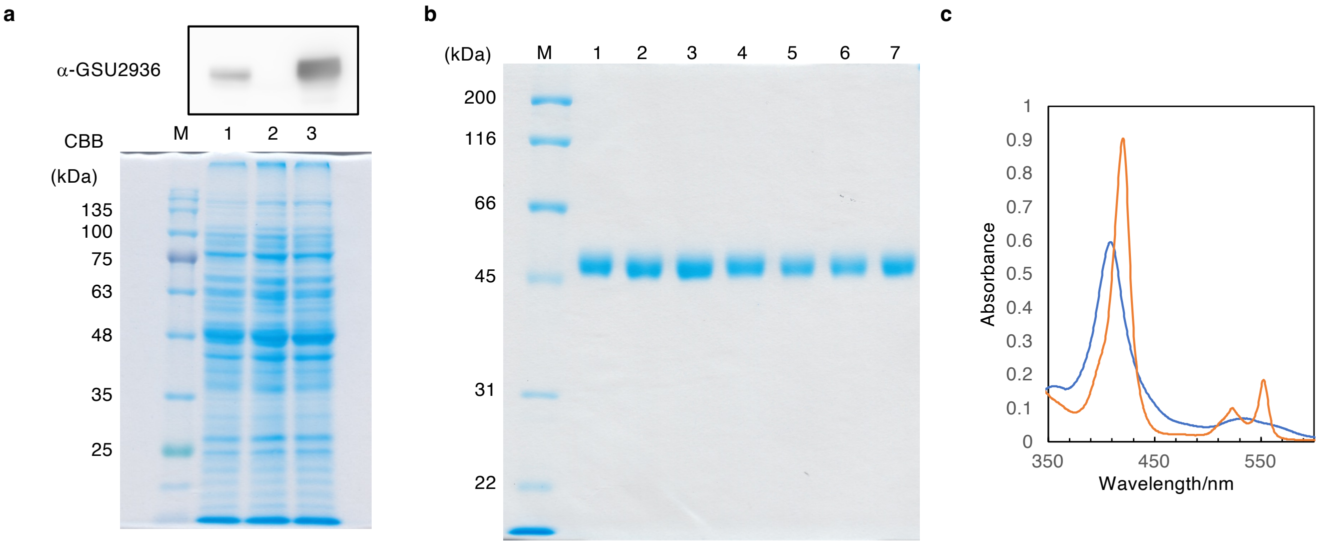


**Extended Data Fig. 3 Expression and purification of recombinant MccSep.**

**a,** Western blot analysis of the crude extracts from the parent strain *G. sulfurrreducens* PCA (lane 1), Δ*mccsep* (lane 2), and Δ*mccsep* complemented with pMcasMccSepStag carrying the *mccsep* gene (lane 3). Crude extracts of *G. sulfurreducens* strains grown in NBAYFE medium were separated on 10% SDS-PAGE gels and analyzed by western blotting using anti-GSU2936 antibody (upper panel). The lower panel represents the identical gels visualized by CBB staining to show the amount of proteins loaded. **b,** Purification of recombinant MccSep (lane 1) and its mutants (lanes 2, U325A; 3, U325C; 4, C239H; 5, C240A; 6, R83A; and 7, K372A). Each recombinant enzyme was purified as a Strep-tag fusion protein and analyzed by SDS-PAGE. Proteins were visualized by CBB staining. **c,** UV-vis absorption spectra of air-oxidized (blue line) and 0.02 mM sodium dithionite-reduced (orange line) MccSep (0.06 mg/mL).

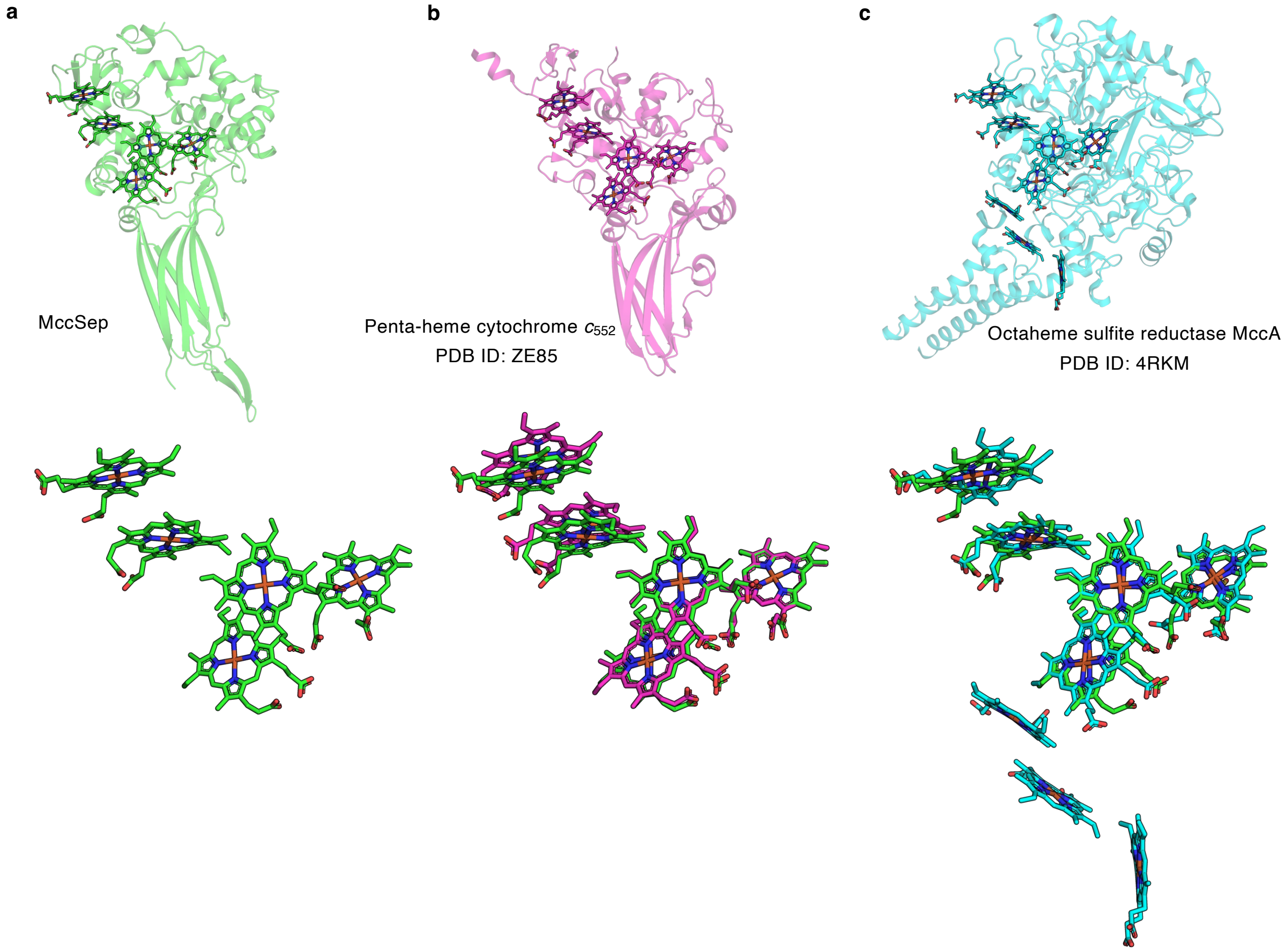


**Extended Data Fig. 4 Comparison with multiheme binding proteins.**
Structural comparison of MccSep with representative multiheme-binding cytochromes. Upper panels show monomeric structures; lower panels display superimposed heme groups aligned with those of MccSep. **a,** MccSep. **b,** Penta-heme cytochrome $c_{552}$ from *Thermochromatium tepidum* (PDB ID: ZE85). **c,** Octaheme sulfite reductase MccA from *Wolinella succinogenes* (PDB ID: 4RKM).

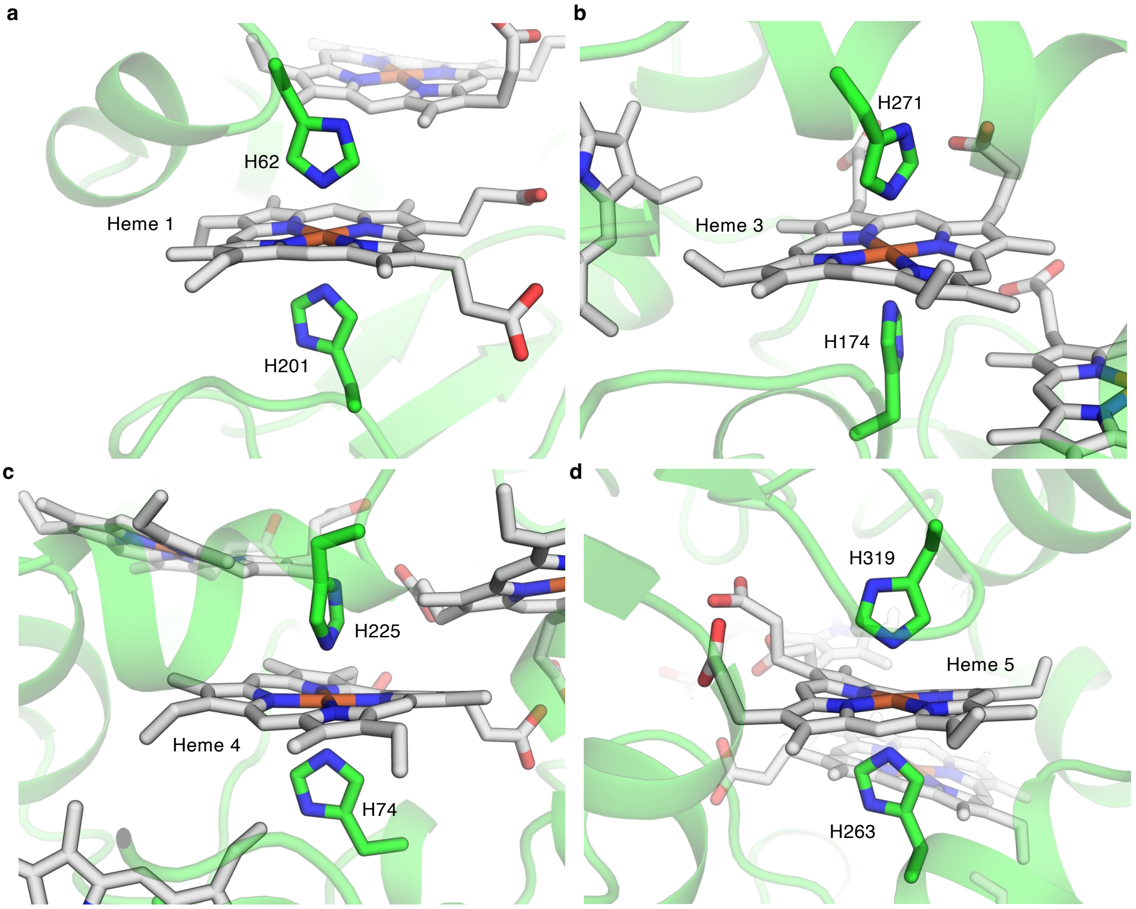


**Extended Data Fig. 5 Heme binding sites in MccSep.**
Each of heme 1 (**a**), heme 3 (**b**), heme 4 (**c**), and heme 5 (**d**) is coordinated by two histidine residues, forming a bis-histidine axial ligation.

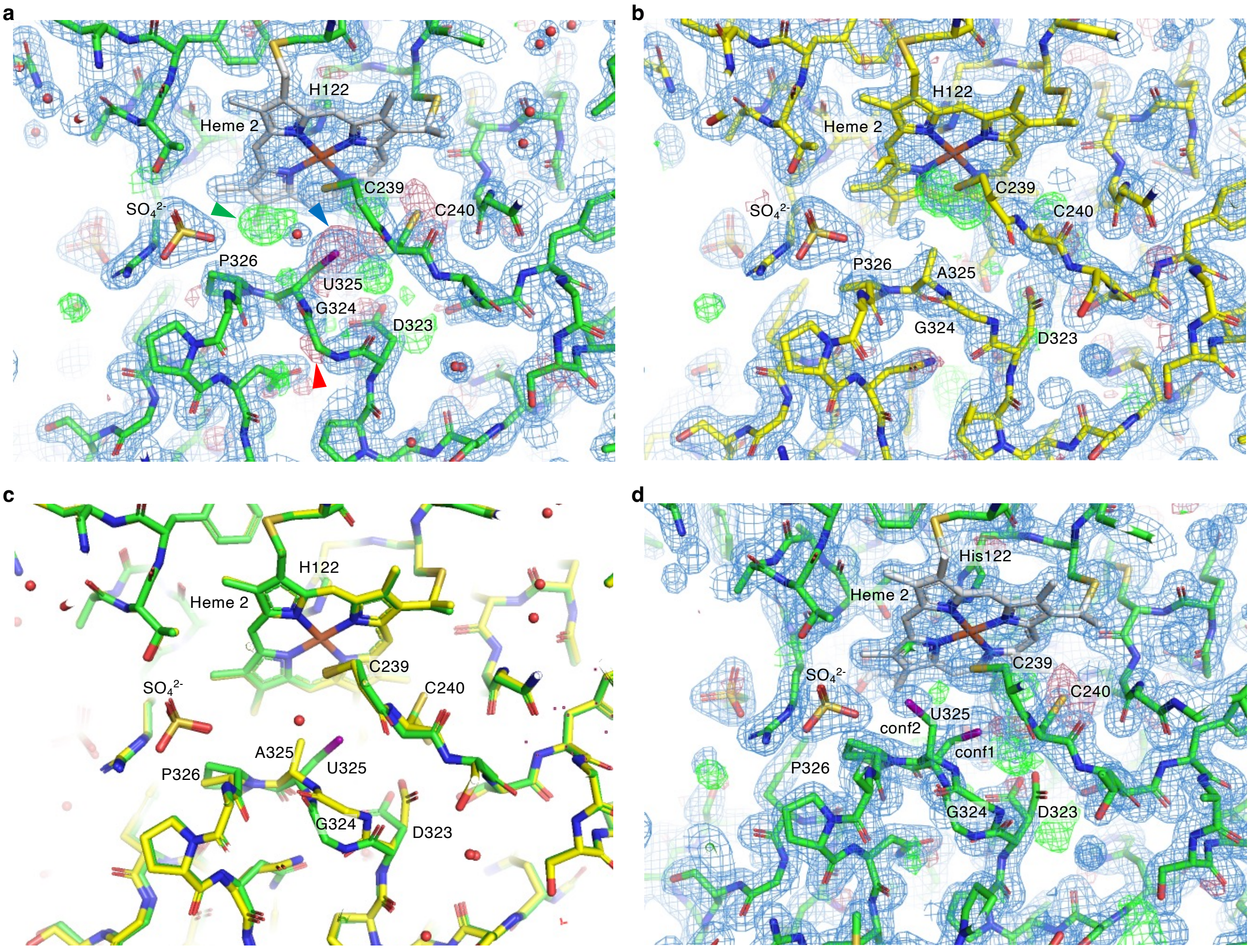


**Extended Data Fig. 6 Assignment of alternative conformations of the 323-326 loop.**
**a,** Initial refinement of the WT MccSep structure showing ambiguous electron density near U325. Negative $F_o - F_c$ electron density peaks are shown on the main chain of G324 (red arrowhead) and the side chain of U325 (blue) with a positive peak ~3.0 Å from the oxygen atom of a sulfate ion (green). **b,** Initial refinement of the U325A mutant showing the loop adopts a single conformation with clear electron density, except for a residual positive peak near. **c,** Superposition of the WT and U325A structures revealing differences in the $C_\alpha$-$C_\beta$ orientation of residue 325, the main chain of G324, and the carbonyl group of D323. **d,** Partially overlapping $2F_o - F_c$ electron densities in the WT MccSep structure with the U325A structure suggest alternative loop conformations in WT. Refinement of the WT structure with dual conformations improves map interpretability. All maps: $2F_o - F_c$ (blue, 1.2σ); $F_o - F_c$ positive (green, 4σ); $F_o - F_c$ negative (red, –3σ).

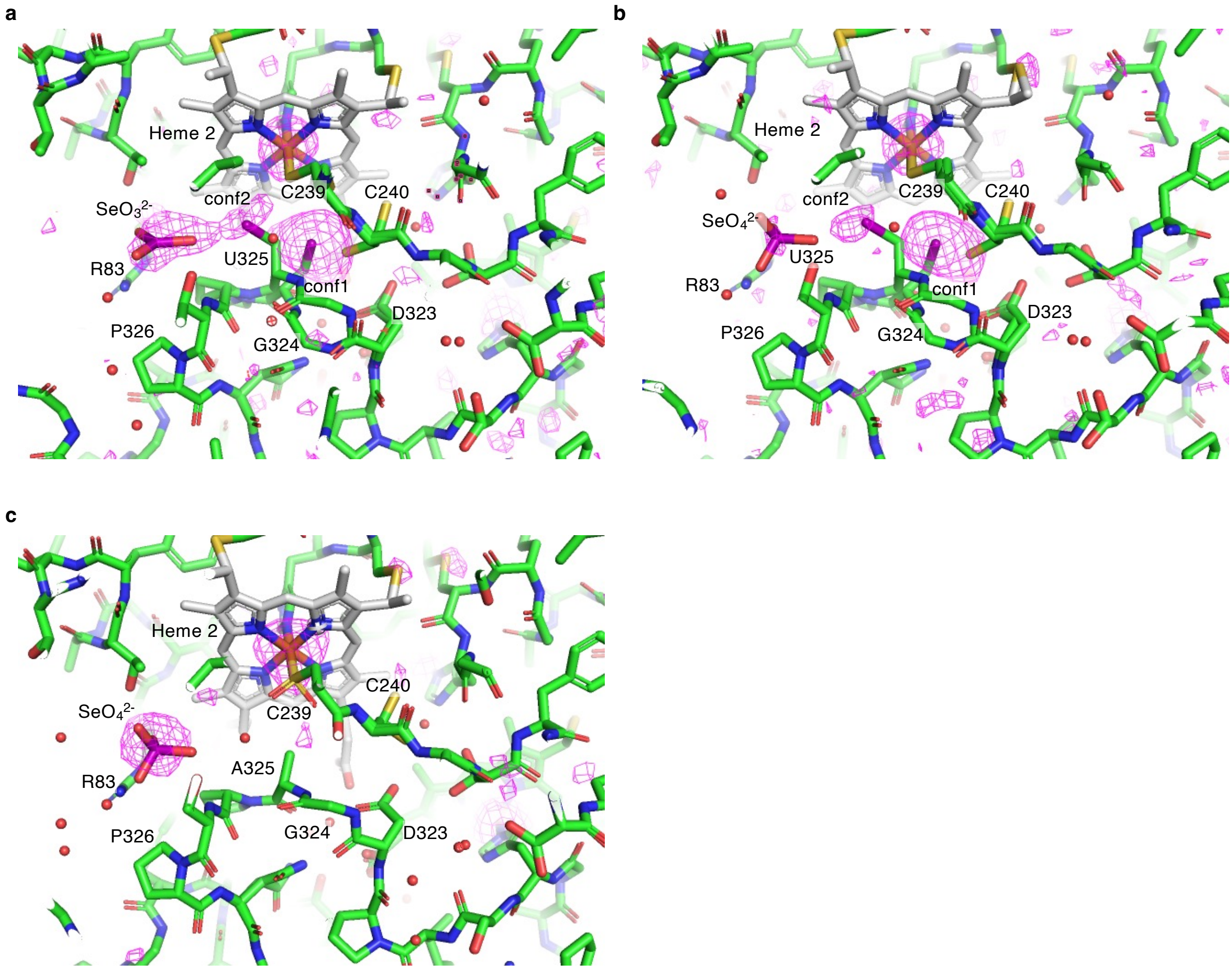


**Extended Data Fig. 7 Anomalous difference Fourier maps at the active site.**
**a–c,** Anomalous difference Fourier maps (magenta, 2.5σ, λ = 0.9792 Å) for WT MccSep soaked with 10 mM selenite overnight (a), 10 mM selenate overnight (b), and for U325A soaked with 100 mM selenate (c). Anomalous difference Fourier map for WT MccSep soaked with 10 mM selenite (a) shows continuous electron density between U325 in conf2 and the bound selenite.
