## Extended Data Tables for "Multiheme selenoenzyme essential for elemental sulfur respiration"

**Extended Data Table 1 Kinetic parameters for polysulfide reduction by MccSep and its mutants**

Data are shown as means ± standard errors of triplicate measurements.

| Enzyme | $V_{max}$ (μmol $MV^{\bullet +}$ $min^{-1}$ $mg^{-1}$) | $K_{0.5}$ (mM) | $h$[a] | $V_{max}/K_{0.5}$ (μmol $MV^{\bullet +}$ $min^{-1}$ $mg^{-1}$ $mM^{-1}$) |
|---|---|---|---|---|
| MccSep | 83 ± 7.0 | 0.19 ± 0.023 | 1.9 ± 0.31 | (4.4 ± 0.66) × $10^2$ |
| R83A | 16 ± 0.78 | 0.21 ± 0.011 | 3.8 ± 0.69 | 73 ± 5.3 |
| C239H | n.d.[b] | n.d.[b] | n.d.[b] | n.d.[b] |
| C240A | 70 ± 4.8 | 0.14 ± 0.015 | 2.1 ± 0.45 | (4.9 ± 0.61) × $10^2$ |
| U325A | n.d.[b] | n.d.[b] | n.d.[b] | n.d.[b] |
| U325C | n.d.[b] | n.d.[b] | n.d.[b] | n.d.[b] |
| K372A | 5.6 ± 0.27 | 0.074 ± 0.0088 | 2.0 ± 0.48 | 76 ± 9.7 |

[a]Hill coefficient

[b]n.d.: not determined due to no detectable activity

**Extended Data Table 2 Data collection, phasing, and refinement statistics**

| | WT native | WT I-derivative | U325A | WT-selenite | WT-selenate | U325A-selenate |
|---|---|---|---|---|---|---|
| PDB accession code | 9WT8 | | 9WT9 | 9WTA | 9WTB | 9WTC |
| **Data collection** | | | | | | |
| Space group | $P2_1$ | $P2_1$ | $P2_1$ | $P2_1$ | $P2_1$ | $P2_1$ |
| Cell dimensions | | | | | | |
| *a*, *b*, *c* (Å) | 83.03 123.54 116.52 | 83.34, 122.98, 116.76 | 83.17 122.65 116.45 | 83.45 123.56 116.61 | 83.64 123.83 116.62 | 83.39 123.62 116.64 |
| β (°) | 92.19 | 92.28 | 92.15 | 92.15 | 91.94 | 91.60 |
| Resolution (Å) | 48.54 - 1.75 (1.81 - 1.75) * | 50.00 - 2.40 (2.46 - 2.40) * | 48.53 - 1.80 (1.86 - 1.80) * | 45.25 - 2.20 (2.28 - 2.20) * | 46.10 - 2.20 (2.28 - 2.20) * | 42.41 - 2.20 (2.28 - 2.20) * |
| $R_{sym}$ or $R_{merge}$ | 0.063 (0.981) | 0.061 (0.981) | 0.081 (1.931) | 0.081 (0.934) | 0.083 (0.810) | 0.072 (0.899) |
| $I / \sigma I$ | 18.12 (2.03) | 21.75 (6.85) | 14.33 (0.90) | 17.58 (2.20) | 16.45 (2.38) | 19.89 (2.32) |
| Completeness (%) | 99.3(98.6) | 95.1 (91.8) | 99.8 (99.7) | 99.9 (100.0) | 99.9 (100.0) | 99.9 (100.0) |
| Redundancy | 7.1 (7.4) | 14.6 (15.0) | 7.0 (7.3) | 6.9 (7.1) | 6.9 (7.1) | 6.9 (7.1) |
| **Refinement** | | | | | | |
| Resolution (Å) | 48.54 - 1.75 (1.81 - 1.75) * | | 48.53 - 1.80 (1.86 - 1.80) * | 45.25 - 2.20 (2.28 - 2.20) * | 46.10 - 2.20 (2.28 - 2.20) * | 42.41 - 2.20 (2.28 - 2.20) * |
| No. reflections | 234,125 (23,146) | | 215,139 (21,395) | 119,604 (11,857) | 120,174 (11,920) | 119,673 (11,904) |
| $R_{work}$ / $R_{free}$ | 0.1669 (0.2865) / 0.1981 (0.3149) | | 0.1870 (0.3828) / 0.2243 (0.4084) | 0.1646 (0.2823) / 0.2066 (0.3186) | 0.1644 (0.2654) / 0.2104 (0.3000) | 0.1682 (0.2846) / 0.2140 (0.3255) |
| No. atoms | | | | | | |
| Protein | 13,426 | | 13,218 | 13,429 | 13,445 | 13,265 |
| Ligand/ion | 1,458 | | 1,181 | 1,224 | 1,195 | 1,063 |
| Water | 1,141 | | 1,038 | 962 | 949 | 916 |
| *B*-factors | | | | | | |
| Protein | 33.27 | | 39.89 | 46.38 | 44.82 | 46.11 |
| Ligand/ion | 34.98 | | 39.78 | 46.51 | 44.26 | 43.98 |
| Water | 41.85 | | 44.87 | 49.99 | 48.46 | 46.36 |
| R.m.s deviations | | | | | | |
| Bond lengths (Å) | 0.009 | | 0.009 | 0.009 | 0.009 | 0.009 |
| Bond angles (°) | 1.05 | | 1.06 | 0.96 | 0.97 | 0.98 |

*Each dataset was collected from one crystal. *Values in parentheses are for highest-resolution shell.
