## Supplementary Information for "Multiheme selenoenzyme essential for elemental sulfur respiration"

or

Hiroyoshi Matsumura, College of Life Sciences, Ritsumeikan University, Kusatsu, Shiga 525-8577, Japan.

**Supplementary Table 1. Oligonucleotides used**

| Primer | Sequence (5´ to 3´) |
|---|---|
| gsu2935-F | AATGCTACGGCTGTCATACGAAATA |
| gsu2935-R | TTTCCCCTTGAAGGTAGAGACGTAG |
| 16SrRNA-F | TGAGACACGGTCCAGACTCCTAC |
| 16SrRNA-R | TCA TTTCTTCCCTCCCGACA |
| GSU01 | AGAAGGGCGCCCATGTTC |
| GSU02 | TGATCCCCGTATTGGTAT |
| GSU03 | TCGATCCATGCCCGCTCAAGGATGAATGTCAGCTAC |
| GSU04 | TCCACTTCCAGATCGTGGAGAAGGCGGCGGTGGAATCG |
| GSU05 | CCACGATCTGGAAGTGGA |
| GSU06 | TGAGCGGGCATGGATCGA |
| GSU3219F | GGAAGGCACGTACAACGGAGTCAAGAGC |
| GSU5417R | GCACCCGGCAGCACTAAGGAACATAACG |
| GSU2936-F | GGAATTCCATATGCCGACCCCCAACCGACTGGTTCTG |
| GSU2936-R | CTCCGCTCGAGCTTGCCCTTGGCGCTGATG |
| casF | AATTGACTGACTGAGAAGAAGATATACATATGCCCTCGAGTCTAGAG |
| casR | AATTCTCTAGACTCGAGGGCATATGTATATCTCCTTCTCAGTCAGTC |
| MccSep-F | ATGGATCCCATATGAAGCACAAAACCGCATG |
| MccSep-Stag-R | TTTTTCGAACTGCGGGTGGCTCCACCTACCTTCGATCTTGCCCTTGGCGCT |
| Stag-Xba-R | AATTTCTAGATTATTTTTCGAACTGCGGGTGGCTCCA |
| MccSep-R | AATTTCTAGACTACTTGCCCTTGGCGCTGATGC |
| C240A-F | CCGACCCAGTGCGCCACCAGCTATGCGAGC |
| C240A-R | GCTCGCATAGCTGGTGGCGCACTGGGTCGG |
| C325A-F | ATCCCCGATGGCTGCCCGACCCCCAACCG |
| C325A-R | CGGTTGGGGGTCGGGCAGCCATCGGGGAT |
| C325C-F | CGATCCCCGATGGCGCCCCGACCCCCAACC |
| C325C-R | GGTTGGGGGTCGGGGCGCCATCGGGGATCG |

**Supplementary Method 1 Reverse transcriptional PCR**

Total RNA extraction from each *G. sulfurreducens* strain was performed using a Sepasol-RNA I Super G kit (Nacalai Tesque). DNase treatment was performed to prevent DNA contamination. RNA quality was assessed by 1% agarose gel electrophoresis, and RNA concentration was determined using absorbance at 260 nm on a Shimadzu UV-1850 spectrophotometer. cDNA synthesis was carried out with a ReverTra Ace qPCR RT Master Mix with gDNA Remover kit (Toyobo) using a DNA Thermocycler (Bio-Rad). PCR reaction containing 200 ng of cDNA, an appropriate primer set, and a KOD Plus Neo PCR kit (Toyobo, Japan) was performed.
